# Non-enzymatic error correction in self-replicators without extraneous energy supply

**DOI:** 10.1101/2025.06.26.661679

**Authors:** Koushik Ghosh, Parthasarthi Sahu, Shashikanta Barik, Hemachander Subramanian

## Abstract

Accurate propagation of sequence information in nucleic acids is central to the evolutionary dynamics of self-replicating systems. Modern biological systems achieve high fidelity using enzymes that actively correct errors through energy-driven mechanisms. However, such complex machinery was absent under prebiotic conditions. Here, we present a theoretical model of error correction in self-replicating heteropolymers that requires neither enzymes nor an extraneous energy supply. The model relies solely on the free-energy gradient driving strand growth and requires asymmetric cooperativity — a kinetic asymmetry known to promote unidirectional elongation. Despite its simplicity, we demonstrate that this minimal model facilitates kinetic discrimination between correct and incorrect base pair incorporations, and for specific set of parameters, reproduces the error ratio of ∼ 10^−4^, experimentally observed in passive base selection processes. It replicates key features observed in DNA error correction, including stalling, fraying, next-nucleotide effects, and the speed–accuracy trade-off. Our results provide plausible answers for longstanding questions, such as the energy source for the enhanced base selectivity of passive DNA polymerases and the role of thermodynamics and kinetics of phosphodiester bond formation in error correction. We show that catalysis of the phosphodiester bond plays a central role in error correction, even without explicit enzymatic structural discrimination. This observation points to a plausible pathway for accurate oligomer synthesis under prebiotic conditions, driven solely by the thermodynamic gradient favoring strand elongation. More broadly, the model highlights how persistent molecular order can emerge from non-equilibrium dynamics – a central requirement for emergence of life.

## 1. Introduction

Heritable genetic information and the occasional alteration of this information through mutations form the bedrock over which evolutionary trajectories of organisms are carved out [1, 2]. In modern cellular systems, the error rates of biochemical reactions are significantly lowered by an energy-driven enzymatic mechanism known as “kinetic proofreading” [3, 4]. This process utilizes external energy source, typically the hydrolysis of energy-rich molecules such as ATP or GTP, to facilitate reaction cycles that enhance accuracy, albeit with increased energy consumption. In the primordial scenario, heteropolymers that self-replicated non-enzymatically possibly accumulated mutations at an unsustainable rate to carry information (probably pertaining to autocatalysis) across generations. Such high mutation rates preclude the possibility of the emergence of error-correcting enzymes, leading to the classic chicken-and-egg problem. How did the primordial molecular evolution escape this fate?

In this work, we provide a possible answer to this question by introducing a theoretical model of non-enzymatic self-replication that relies solely on *thermodynamic drive* – the net free energy change favoring templated daughter strand formation over dissociation, which in biochemical contexts can arise from incoming nucleotides or environmental conditions [5–7]. Below, we review existing approaches to this problem and discuss their limitations, before presenting our model and the insights it provides on non-enzymatic error correction.

### i. The hypercycle approach

Manfred Eigen proposed a mechanism to evade the decay of information under high mutation rates by assuming an autocatalytically closed set of self-replicators, called quasispecies, that replicated with high fidelity, as long as the mutation rate was below a certain threshold [8, 9]. This proposal allowed for the possibility of stable information transmission across generations of quasispecies, as long as the error rates are below a threshold that depends inversely on the heteropolymer length. However, it is difficult to imagine an evolutionary pathway from this hypercyclically coupled community-based error tolerance mechanism to the emergence of a single-molecule-based information storage and error correction system. This is because the hypercyclic coupling leads to an evolutionary preference for multiple error-prone, catalytically coupled self-replicators over a single, more accurate, uncoupled self-replicator. While the quasispecies model has provided valuable insights into the highly error-prone viral evolution [10], especially for RNA viruses, its applicability to the evolution of stable information transfer in primordial non-viral systems is less clear. This leads us again to the following question: How did single-molecule self-replicators develop an ability to correct errors without enzymatic/community help?

### ii. Kinetic Proofreading

Although extraneous energy-driven, it is important to discuss the paradigmatic Hopfield-Ninio scheme of enzymatic proofreading [3, 4] here, to distinguish our model from the above scheme. Hopfield-Ninio model involves a reaction topology in which at least two discriminatory dissociation paths are available to differentiate between the correct and incorrect substrates. When both of these discriminatory paths are optimally utilized by driving the reaction flow using an external thermodynamic drive, maximal error correction, that is better than the thermodynamic limit, is achieved. The thermodynamic drive, usually the hydrolysis of ATP, drives the reaction towards not just product formation, but also towards an additional dissociation path for both the correct and incorrect substrates, resulting in the expenditure of energy even for futile reaction cycles that do not result in product formation. This model has been experimentally verified by observing this excess ATP consumption during translation soon after the model was introduced [11]. Because the accuracy of the substrate selection improves when the incorrect substrate cycles through the reaction a large number of times, the corresponding energy expenditure for maximal error correction becomes significantly higher, particularly in the presence of high incorrect substrate concentrations [11–14]. In addition, the rate of product formation must also be low to allow the incorrect substrate to cycle through the reaction many times, thereby introducing a time lag. This requirement in favor of accuracy reduces the rate of product yield, and results in a speed-accuracy trade-off, with the maximal error correction occurring when the product yield rate tends to zero. More complex reaction topologies that are derived from the above simple scheme inherit the same issues, although regimes in the parameter space have been found where the speed is improved significantly for a tolerable loss of accuracy[12, 15, 16]. This suggests that we need models that maximize self-replication accuracy at non-zero reaction speeds, without requiring extraneous energy sources for driving the reaction and for error correction.

### iii. Base selection energetics

Error correction in DNA replication is a multistage process, beginning with the stage of *base selection*, wherein the polymerase *passively* selects the correct bases over incorrect ones [17], without the expenditure of additional energy. The energetic foundation of fidelity during this initial phase of *in vivo* DNA replication remains incompletely understood [17–23]. Subsequent stages involve exonuclease-mediated proofreading and post-replicative mismatch repair, which, however, are not strictly necessary for cell survival [23, 24]. During the base selection stage, the error rate decreases from approximately 10^−2^, determined thermodynamically by the free energy difference between correct and incorrect base pairs in an aqueous medium, to about 10^−4^, as observed in exonuclease-deficient polymerase alpha and Taq polymerases [17, 23].

One proposed explanation for this enhanced selectivity is that the polymerase active site excludes water molecules, thereby increasing the enthalpic discrimination between correct and incorrect base pairs [25, 26]. This explanation, however, appears to challenge Landauer’s principle since it suggests improvement in accuracy without concomitant energy expenditure, allowing the enzyme to function as a Maxwell’s Demon [27]. For instance, during the replication of a homonucleotide DNA template, in the absence of the polymerase, the replication would result in a disordered daughter strand with a considerable number of random point mutations, whereas, in the presence of the passive polymerase, a more ordered, low entropy daughter strand with fewer errors is produced, apparently without any concomitant extraneous energy expenditure. Conceptually, if two such reactions are carried out to equilibrium — one with the polymerase and one without — and their chemically distinct products are mixed, the resulting free energy gradient could, in principle, be harnessed to perform work. This would violate the first law of thermodynamics, suggesting that passive base selection must involve an unrecognized energetic cost [23]. The model we present below identifies the source of this energetic cost as the one associated with the thermodynamic drive for self-replication itself.

### iv. Other approaches

Here, we briefly review more recent attempts at demonstrating non-enzymatic self-replication and error correction in models of primordial self-replication. Enzyme-free template-directed polymerization has been extensively studied experimentally as a model for primordial self-replication [28, 29]. These studies have highlighted a critical challenge: high error rates — often exceeding 10% — arising from the absence of intrinsic error correction mechanism [30, 31]. In response, various theoretical and experimental approaches have been developed to enhance fidelity under prebiotic conditions [32–34].

One such approach, known as kinetic error filtering [32], builds on the observation that primer extension slows down the incorporation of a mismatch. This slowdown is included in the theoretical model to reduce the replication of erroneous strands, which can then be selectively removed by applying a limited time window for self-replication. Another recent model by Matsubara et al.[33] presents a proofreading mechanism based on positive feedback between polymerization kinetics and template population dynamics, aiming to operate under broadly prebiotic conditions without enzymes. While this model offers an intriguing route to error correction, it involves assumptions such as the strong dependence of monomer addition rates on the thermodynamic stability of the primer-template complex and immediate strand separation after extension. These assumptions, along with the observed slow timescales and low yields for longer templates, present challenges for practical applicability — especially for longer, information-carrying strands. A complementary approach by Mukherjee et al.[34] implements non-enzymatic error correction experimentally using DNA strand displacement networks fueled by chemical energy. This design effectively mimics the Hopfield-Ninio proofreading scheme, enhancing discrimination through multiple reversible binding events and energy-driven cycling. Another mechanism proposed in [35], where spatial separation between substrate binding and product formation events is used to impose time delays that prevent wrong substrates from reaching the product formation site, improving accuracy rates. However, even this setup requires the maintenance of concentration gradients — a condition that likely demands external energy input. While each of these strategies contributes valuable insight into potential prebiotic error correction mechanisms, many rely predominantly on static thermodynamic considerations or require organized energy inputs. These requirements may pose challenges for their implementation in prebiotic conditions, where such organized energy inputs were likely unavailable [36–38].

### v. Persistent Order from Kinetics

More generally, the search for a model system that autonomously creates order from non-equilibrium supply of nutrients and energy, that persists on a timescale longer than the timescale of the energy supply, is still ongoing [39–42]. Demonstrating that such a system can be spontaneously created in the primordial scenario, *de novo*, would provide us confidence in our belief of spontaneous emergence of life on early Earth. It would also enable us to investigate and understand the physicochemical mechanisms for creating persistent order from free energy flow and implement them in self-repairing biomimetic systems. Our proposed model illustrates a potential kinetic mechanism for the emergence of persistent order in non-enzymatic self-replicating systems under plausible prebiotic conditions. In equilibrium systems like magnets, broken symmetry leads to long-range spatial order. Similarly, in our non-equilibrium model, broken symmetry gives rise to persistent order over time, maintained through error correction.

In the following, we present our theoretical model showing that non-enzymatic self-replication can achieve high fidelity by utilizing the thermodynamic drive for self-replication itself, without any extraneous energy input. We also address the conceptual challenges discussed in the above paragraphs. The central ingredient of our model is *asymmetric cooperativity* — a kinetic property previously introduced to understand the counterintuitive evolutionary choice of unidirectional replication of DNA strands on a template [43, 44]. We adopt the existence of asymmetric cooperativity in our system as the *central premise* of this article, and explore its implications for the fidelity of self-replication.

Importantly, our model shows that the ability to correct errors depends critically on the unidirectional growth of the daughter strand; disrupting this directionality results in a loss of self-replication fidelity. Within this model, we demonstrate that the error rate can be reduced from the thermodynamic baseline of approximately 10^−2^ to around 10^−4^ — approaching the fidelity observed in passive polymerases — without the need for any external energy input. Moreover, the model naturally reproduces several experimental features commonly associated with error correction: (a) stalling, where strand elongation pauses following the incorporation of an incorrect base pair, consistent with observations in non-enzymatic DNA replication [45–47]; (b) a speed-accuracy trade-off, analogous to trends seen in enzymatic proofreading systems [48–51]; (c) fraying or unzipping of the primer terminus upon incorrect base pairing at the elongation site [52]; (d) next-nucleotide effects [53], in which the addition of a correct nucleotide after an incorrect one stabilizes the mismatch and elevates local error rates [46]; and (e) adaptive modulation of error rates, as observed in organisms subjected to nutrient stress [54, 55].

We define the thermodynamic drive as the free energy change favoring base pairing and covalent bond formation between nucleotides, under conditions that promote duplex stability, such as temperatures below the dsDNA-to-ssDNA melting transition and/or increased nucleotide concentrations. We consider the covalent bond formation to be effectively irreversible due to its highly exergonic nature (≈ −12*k*_*B*_*T*)[56]. It is also important to note that the present model is intended to capture only the base selection dynamics of self-replication; more advanced mechanisms, including exonuclease-assisted proofreading and mismatch repair, lie beyond the scope of this study.

## 2. Asymmetric Cooperativity Model

For completeness, we provide a brief summary of the asymmetric cooperativity model, introduced in[43, 44]. Asymmetric cooperativity is defined as an asymmetric *kinetic* influence of a pre-existing base pair (hydrogen bond) between a lone nucleotide and a template strand on the formation or dissociation of neighboring base pairs between other nucleotides and the template, to its left and right. Specifically, in the model, we assume that a base pair between a lone nucleotide and the template strand reduces the kinetic barrier for the formation/dissociation of a new base pair towards the 5^*′*^-end of the template, and increases the kinetic barrier for formation/dissociation of a base pair towards the 3^*′*^-end. This kinetic property helps to simultaneously satisfy two conflicting requirements for rapid daughter strand construction: a low kinetic barrier for the induction and easy incorporation of nucleotides to the base pair’s 5^*′*^-end, thereby enhancing the base pairing propensity with a template nucleotide, and a high kinetic barrier to the 3^*′*^-end to retain the base-paired nucleotides on the template strand, facilitating the formation of intra-strand covalent bonds that extend the nascent daughter strand[43]. This directional asymmetry in the kinetic influence of a base pair on its left and right neighbors results in the unidirectional daughter strand construction, and has been shown to improve the rate of self-replication of template [43, 44]. This phenomenon is illustrated in Fig.1. In [44], we termed this as sequence-*independent* asymmetric cooperativity. Here, we simply call it asymmetric cooperativity, since in this article there is no need to invoke sequence-*dependent* asymmetric cooperativity. *The central premise of this article is that the above-mentioned asymmetry cooperativity, a form of kinetic asymmetry [57], is present in correct (complementary) base pairing, whereas it is absent in incorrect (non-complementary) base pairing*. To be precise, we assume that complementary, correct base pairs reduce the kinetic barrier of the 5^*′*^-end neighbor and increase the barrier of the 3^*′*^-end neighbor, whereas incorrect base pairs neither reduce nor increase the kinetic barriers for base pair formation/dissociation in either direction.

**Figure 1:**
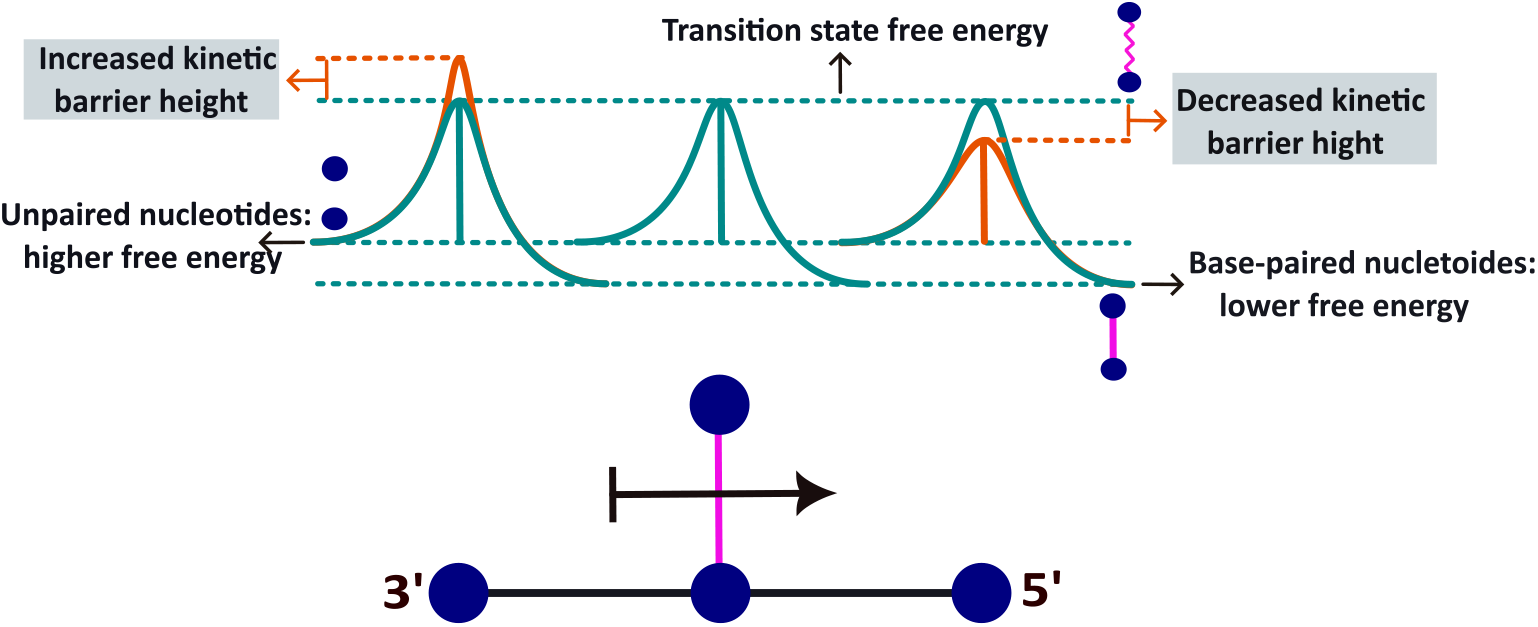
Schematic illustration of asymmetric cooperativity. Nucleotides are denoted by blue circles, and vertical lines between nucleotides represent hydrogen bonds. The associated energy diagram for base pairings is presented above the illustration. The presence of a base pair lowers the kinetic barrier of its right neighboring base pair, denoted by an arrowhead, while increasing the barrier of the left neighbor, denoted by a barhead. This results in favorable conditions for new base pair formation to the right and increased intra-strand covalent bond formation probability to the left due to the increased kinetic barrier that stabilizes the left base pair. This kinetic asymmetry thus establishes efficient unidirectional daughter strand construction by breaking the left-right symmetry.

The mechanism of error correction within our model can be succinctly explained as follows: Let us assume that an incorrect base pair forms at the location *m*, as shown in Fig.2. According to the central premise of our article, this base pair does not influence the kinetic barrier of the already formed base pair at the (*m* − 1)^*th*^ location or for the formation of a new base pair at the (*m* + 1)^*th*^ location. Therefore, the growth of the new daughter strand is stalled, as the barrier towards the template’s 5^*′*^-end is not reduced. This provides more time for the incorrect base pair to dissociate, which is also catalyzed by the reduced barrier at the *m*^*th*^ location due to the catalytic effect of the base pair at the (*m* − 1)^*th*^ location. Moreover, since the base pair at the (*m* − 1)^*th*^ location is not stabilized due to the reduction in *its* kinetic barrier caused by the (*m* − 2)^*nd*^ base pair and the lack of inhibition from the *m*^*th*^ incorrect base pair, it is also prone to dissociate, which can precipitate sequential unzipping of the rest of the bonded segment due to reduced barriers at the Y-fork. This stalling and unzipping behind the growth front significantly delays the strand construction process. This time delay between the correctly and incorrectly formed base pairs can be utilized for error correction if the rate of incorporation of the correct base pair in the growing daughter strand through intrastrand covalent bond formation is fast enough. At very low covalent bond formation rates, the ratio between the probability of the covalent bond formation incorporating the wrong base pair into the growing strand and the probability of the covalent bond incorporating a correct base pair, are nearly equal, since at the long times required for the slow covalent bonds to form, the time lag between the correct and incorrect strand formation vanishes. Only when the rate of covalent bond formation approaches either of the two base pairing rates, the probabilities of correct and wrong base pair stabilization through covalent bonding become distinguishable, and error correction, possible. This idea is schematically illustrated in Fig.5. Since the covalent bond formation is a thermodynamically nearly irreversible event [58, 59], if its rate of formation is just as fast or faster than the formation rate of a stretch of cooperatively coupled correct base pairs, the covalent bond kinetics will ensure that only correct base pairs are irreversibly incorporated into the daughter strand. Thus, the catalysis of covalent bond formation is crucial for enhancing the discrimination between correct and incorrect base pairs during daughter-strand construction, without the need for extraneous energy supply. This may also explain the improved base-selection accuracy that happens apparently without any extraneous energy expenditure.

**Figure 2:**
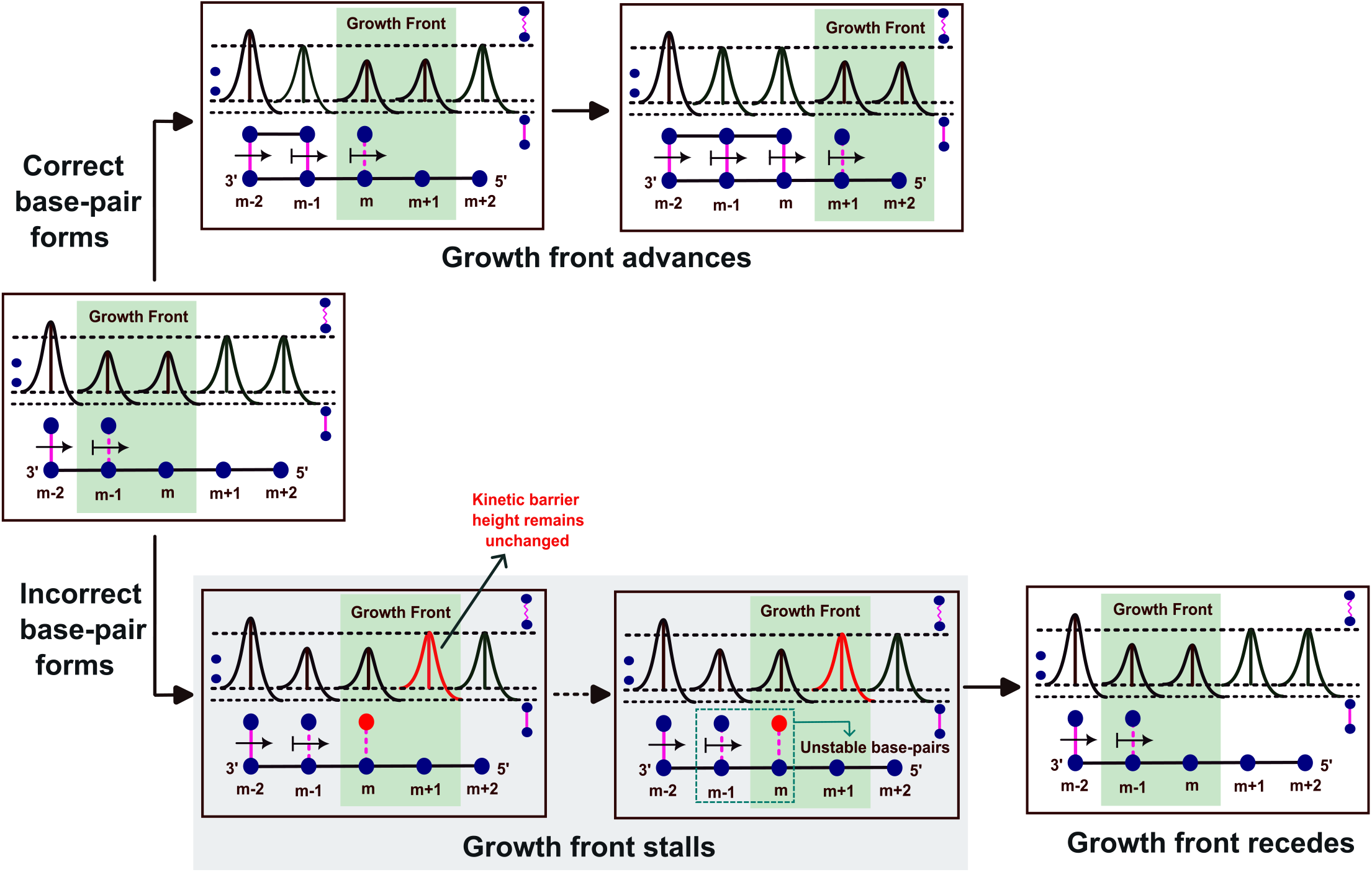
Illustration of error correction during daughter strand construction on a template. Each row shows the pathway for forming a correct (top) or incorrect (bottom) base pair at position *m* on the template, marked by an index below each panel. The three horizontal dotted lines in the energy barrier diagram for base pairing denote energy levels for the bonded (bottom), unbonded (middle) and the transition states (top). The non-complementary nucleotide is represented by a red circle in the bottom panels, which is base paired with the template nucleotide at the *m*^*th*^ location. This incorrect base pair does not catalyze the formation of a new base pair at (*m* + 1)^*th*^ location to its 5^*′*^-end (right), nor does it hinder the dissociation of the (*m* − 1)^*th*^ base pair to the 3^*′*^-end (left), in accordance with the model premises. This can be seen by comparing the barrier heights of these base pairs at the top and the bottom panels. The kinetic barrier of the (*m* − 1)^*th*^ base pair is raised due to inhibition from the *m*^*th*^ correct base pair, stabilizing it (top), whereas, the barrier of (*m* − 1)^*th*^ base pair is unaffected by the presence of *m*^*th*^ incorrect base pair, leaving it unstable (bottom). Similarly, the kinetic barrier of the (*m* + 1)^*th*^ location is lowered due to the presence of the *m*^*th*^ correct base pair, catalyzing the formation of a new base pair and extending the daughter strand (top), whereas, the barrier of the (*m* + 1)^*th*^ location is unaffected by the presence of an incorrect base pair at *m*, making daughter strand extension harder, which also leaves the incorrect base pair unstable, due to its lowered kinetic barrier. The growth front advances in case of correct base pair formation at *m* (top), whereas, the growth front recedes in case of an incorrect base pair (bottom). This results in a time lag between the formation of correct and incorrect base pairs. If the covalent bond formation time is of the same order of the time lag, the irreversible covalent bond formation event can discriminate between the correct and incorrect base pairs, and would lead to error correction. The top panels show the covalent bond forming between the first and second daughter strand nucleotides, whereas no covalent bonds form between these two nucleotides in the bottom panels, due to the second base pair’s instability, illustrated by its lowered kinetic barrier. Thus, covalent bond catalysis leads to error correction.

## 3. Methods

We consider a linear template strand comprising of *N* nucleotides. The initiation of daughter strand construction occurs when one of the free-floating nucleotides forms a base pair with a template nucleotide. An illustration of the daughter strand construction process under the influence of asymmetrically cooperative base pairing is shown in Fig.2. We model the daughter strand construction as a series of base pair formation/dissociation events between nucleotides on a template strand and free-floating nucleotides. Mathematically, these series of events can be modeled as a continuous time Markov chain [60–63] that describes each of these bonding/dissociation events as a transition between any two predefined states. These states are denoted by an *N*-length numerical sequence whose digits can take any of the values from the set {0,1,2}. A value of 0 at any location represents the absence of a base pair between the template nucleotide and another nucleotide at that location, and a value of 1(2) represents the presence of a correct (incorrect) base pair at that location. For example, for a given template strand consisting of *N* = 5 nucleotides, the state 00000 would signify a single template strand with no base pairs present, whereas 11111 and 11211 represent states in which all nucleotides in the template strand are hydrogen-bonded to free-floating nucleotides, either correct or incorrect. To keep our analysis simpler and the size of the state space small, *we allow for an incorrect base pair formation only at a single predetermined location in the strand*.

### I. Calculation of transition rate matrix

The transition rates among these states can be calculated using the base pair formation rate *p* between free-floating and template nucleotides, along with two distinct dissociation rates: *q*_*r*_ for correct base pairs and *q*_*w*_ for incorrect base pairs. To incorporate asymmetric cooperative interactions between neighboring base pairs on the template, we introduce two asymmetric cooperativity parameters *α* and 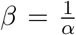 that modify the formation/dissociation rates of the nearest-neighbor base pairs to the right (template’s 5^*′*^-end direction) and left (template’s 3^*′*^-end direction) of a pre-existing base pair, respectively. Fig.3 illustrates the calculation of transition rates between different states used to model the daughter strand construction on a four-nucleotide long linear template strand. The choice of four nucleotides is only for ease of illustration; the computational experiments in the manuscript are performed on templates five nucleotides long. Supplementary material (Section VI) shows that the model is not length-specific, and works for other template lengths as well.

**Figure 3:**
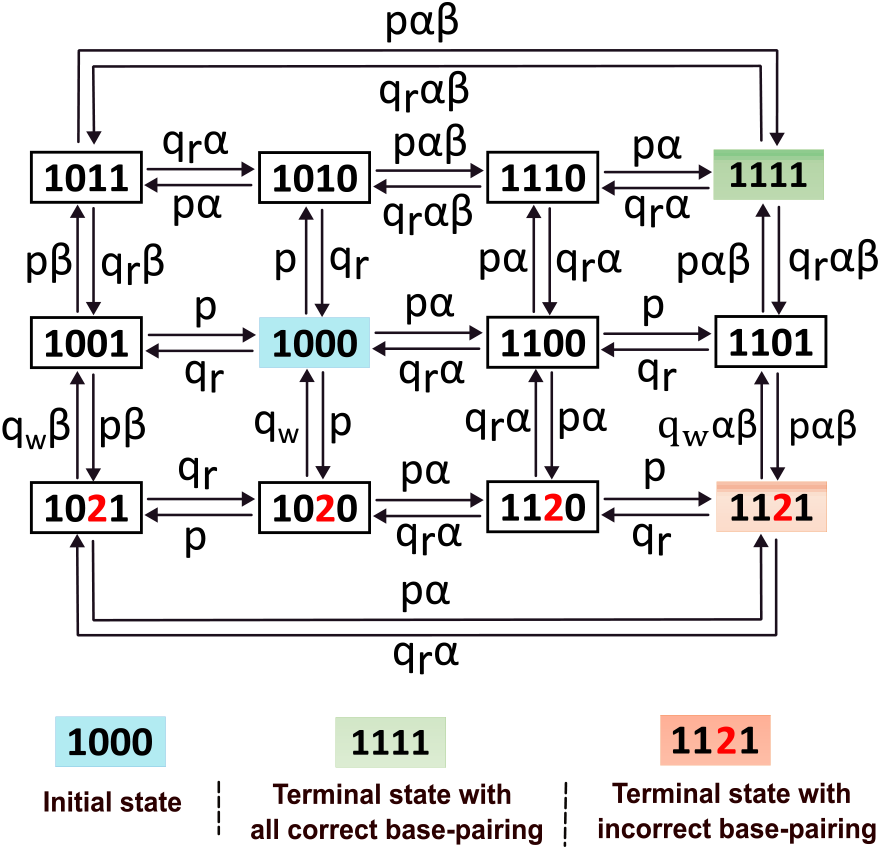
Illustration of calculation of the transition rates between states representing the growth of the daughter strand on a four-nucleotide-long linear template strand, where the first nucleotide plays the role of a primer, and is therefore dynamically inactive. The states are represented by a four-digit-long sequence containing 0, 1, or 2, with 0 denoting the absence of a base pair between the template nucleotide and a free-floating nucleotide, and 1 (2) denoting the presence of a base pair between the template strand nucleotide and a complementary (non-complementary) nucleotide, respectively. The starting state, 1000, represents a single template strand with one base pair at the 3^*′*^-end, whereas, the final state, 1111 (1121), corresponds to all nucleotides of the template strand base paired with complementary (non-complementary at third position) free-floating nucleotides. The transition rates between these states are dependent on the presence or absence of adjacent base pairs to the left and right of a given base pair. The bare formation and dissociation rates, denoted by *p* and *q*_*r*(*w*)_, are modulated by the local context. Specifically, for any given location on the template, if only the left neighboring correct base pair is present, it reduces the kinetic barrier for base pair formation or dissociation at that location by a factor *α*. If only the correct right neighboring base pair is present, it increases the kinetic barrier by a factor *β*. When both the neighboring base pairs are present and are correct, the kinetic barrier for base pair formation/dissociation at the given location is simultaneously reduced and enhanced and is therefore multiplied by a factor *αβ*. Incorrect neighboring base pairs do not affect the formation or dissociation rates.

### II. Time evolution of state occupation probability: Absorbing and non-absorbing target states

In order to understand the daughter strand construction process in the presence of asymmetric cooperativity and intuitively understand the origin of the error correction mechanism, we need to evaluate the transition of the Markov chain across each and every state, starting at the initial state 00000, and ending at either a correctly formed daughter strand 11111 or a daughter strand with an error, 11211. We computationally evaluate the Markov chain trajectory from the initial state to the final state in two different ways: (i) as an absorbing Markov chain, where, once the chain reaches the correct/incorrect final state for the first time, it stops and never returns back. This assumption implies that once all the base pairs are formed, the chain terminates, and the base pair hydrogen bonds are not allowed to break. (ii) as a non-absorbing Markov chain, which allows for the chain to leave the final state and re-enter intermediate states. This assumption allows for breaking of base pairs even after all base pairs are formed.

The time evolution of the occupation probability *P*_*i*_(*t*) of the *i*^*th*^ state involves solving the master equation [60–63].

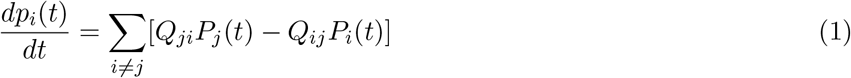

In matrix notation Eq.1 can be written as

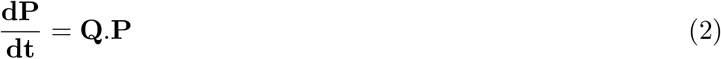

whose formal solution is,

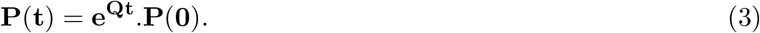

Here, *P* (0) is a column matrix that contains the initial probability for the Markov chain to be in every state at the start of the process, and *P* (*t*) is a column matrix in which each entry *P*_*i*_(*t*) is the probability for the Markov chain to be in the *i*^*th*^ state at time *t*. All elements of *P* (0) are zero, except for the initial state element, which is assigned 1.

For modeling an absorbing Markov chain time evolution, we assign *Q*_*R,i*_ = 0, *i* ≠ *R*, and *Q*_*W,i*_ = 0, *i* ≠ *W*, where *R* and *W* denote the indices of the correct and incorrect final states. This assignment prevents the Markov chain from traversing back to the intermediate states once it reaches the final states. For non-absorbing Markov chain, these rate matrix elements carry their usual values assigned as mentioned above.

We quantify error correction as the ratio, *η*, of the probability of occupation of the correct final state *P*_*R*_ to the probability of occupation of the incorrect final state *P*_*W*_, after the Markov chain reaches steady state, i.e., becomes time-independent, for both absorbing and non-absorbing Markov chains.

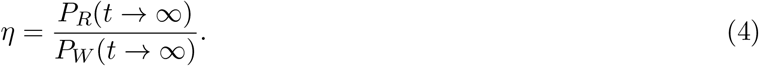

Thus far, we have used the 00000 as the initial state at which the Markov chain begins its sojourn. However, the transition from the state 00000 to any of the single-bonded states (10000, 01000, 00100, 00010, 00001) is independent of the kinetic parameters *α* and *β*, and therefore, sets a kinetics-independent floor value to the mean first passage times for the correct and incorrect final states. This is because of the much-lower rate of the above kinetically-unassisted transition, compared to majority of the state transitions that follow. Since our goal is to demonstrate the necessity of kinetics for error correction, this is counterproductive. Henceforth, we designate the state 10000 as the effective starting point for daughter strand construction in our analysis, and remove 00000 from the state space. This reassignment can be thought of as modeling daughter strand construction through primer extension[64]. The first base pair (primer) is assumed to be dynamically inactive or frozen and is not allowed to break.

### III. Modeling of covalent bond formation

Since the error correction of an incorrect base pair at a location *m* depends on the presence or absence of base pairs at both *m* − 1 and *m* + 1 locations within our model, we impose an experimentally observed [65–67] constraint that the covalent bond formation between the nucleotides in the daughter strand at the locations *m* − 1 and *m* happens *after* the hydrogen bonding of the *m* + 1 base pair. However, at high nucleotide binding rates, the possibility of a base pair forming at *m* + 2 without a base pair at *m* + 1 becomes non-negligible. This configuration hinders the formation of a base pair at *m* + 1, improving the error correction because of the catalysis of the wrong bond dissociation by the bond at *m* − 1. Moreover, a base pair at *m* + 2 stabilizes the base pair at *m* indirectly, by stabilizing *m* + 1, thereby enabling covalent bond formation between *m* and *m* – 1 nucleotides. To include these interactions, we include two base pairs to the left of the wrong base pair location, and two to the right, resulting in a template of length *N* = 5. We impose the constraint that, for a template of length *N* = 5 with a possible incorrect base pair at *m* = 3, the covalent bond would form after the completion of the base pair formation at all 5 locations, that is, after the Markov chain reaches either of the two final states, 11111 or 11211. We model only the covalent bond formation between the 2^*nd*^ and 3^*rd*^ base pairs, and reasonably assume that the covalent bonds between previous nucleotides on the daughter strand have already formed, while the covalent bond between 3^*rd*^ and 4^*th*^, and 4^*th*^ and 5^*th*^ nucleotides are yet to form, and are not included in the model. We also verified that our results considering covalent bond formation between the first two nucleotides of the daughter strand, in addition to the covalent bond formation between the 2^*nd*^ and 3^*rd*^ base pairs, yields qualitatively identical results (see supplementary material (Section I)).

We introduce an intermediate variable *v* to denote the rate of this covalent bond formation, and express it in terms of the rate of hydrogen bond formation of a stretch of base pairs, starting from 10000 to 11111 or 11211. By expressing the covalent bonding rate in terms of hydrogen bond formation rates, we facilitate a quantification of covalent bond catalysis, and later demonstrate that both these rates must be of the same order for effective error correction. For this purpose, we define a dimensionless covalent bonding rate 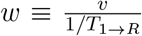, where *T*_1→*R*_ is the conditional mean first-passage time to the fully paired correct final state.

The transition matrix, calculated as illustrated in Fig.3, is inverted numerically, to arrive at the accuracy ratio, Eq.4 and other metrics introduced below. In the supplementary material (Section II), we analytically solve for the time evolution of this system, Eq.1, by introducing certain approximations. The solution is shown to exhibit similar dynamics as the full system, and approximately reproduces the accuracy ratio.

### IV. Stalling: Evaluation of Conditional Mean First Passage Time

To demonstrate the stalling of strand construction due to the formation of an incorrect base pair, we need to evaluate the time taken for the completion of strand construction. This is done by calculating the conditional mean first passage time [60, 68]. The conditional mean first passage time, denoted as *T*_*i*→*k*_, represents the expected time it takes for a system, starting from transient state *i*, to reach absorbing state *k*, given that it eventually reaches *k*. In a continuous-time absorbing Markov chain, the *conditional mean first passage time* from a transient state *i* to an absorbing state *k* is given by

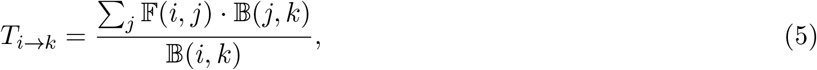

where *F* is the *fundamental matrix*, and *B* is the matrix of *absorption probabilities*. The element *F*(*i, j*) represents the expected total time the process spends in transient state *j* given that it started in transient state *i*, while *B*(*i, k*) denotes the probability that the process, starting from state *i*, is eventually absorbed in absorbing state *k*.

To compute these quantities, we first partition the infinitesimal generator matrix *Q* of the Markov chain as follows:

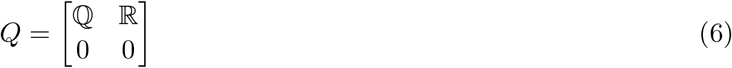

where ℚ is the submatrix corresponding to transitions among transient states, and ℝ contains transition rates from transient states to absorbing states. The fundamental matrix is then given by

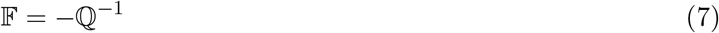

and the matrix of absorption probabilities is computed as

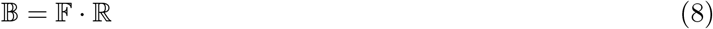

Each row of *B* corresponds to a transient starting state, and each column corresponds to a distinct absorbing state.

Using Eq.5, we calculate the conditional mean first passage time for two possible outcomes of strand construction: *T*_1→*R*_, the expected time to reach the correctly formed daughter strand and *T*_1→*W*_, the expected time to reach a strand containing an error. We then define the *stall quotient τ* as:

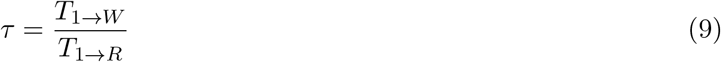

In this study, we examine the dependence of accuracy ratio *η* and stall quotient *τ* on the effective base-pairing drive, defined as the free-energy change associated with correct base-pair formation,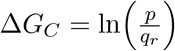 (in units of *k*_*B*_*T*). This quantity reflects only the contribution from correct base pair formation and dissociation and thus represents one component of the total thermodynamic drive for daughter strand construction. In addition, we analyze the influence of the kinetic parameters *α, β* = 1*/α*, and the covalent-bond formation rate *v*, on *η* and *τ*, which collectively determine the kinetics of daughter-strand extension.

### V. Parametrization

#### i. Thermodynamic variables

The thermodynamic contribution to daughter-strand growth arises from the base-pair formation and dissociation rates, denoted *p* and *q*_*r/w*_, which depend on temperature, monomer concentration, ionic strength, and other environmental variables [6]. The effect base pairing drive Δ*G*_*C*_, introduced above, modulates the base-pair association rate and thereby influences the overall strand extension process. It should be noted that, Δ*G*_*C*_, does not represent the total thermodynamic drive for daughter-strand construction. The total thermodynamic drive – which includes all nonequilibrium contributions across the entire strand-construction cycle – is quantified separately through the total entropy production rate under nonequilibrium steady-state (NESS) conditions [69], as described in section III of the supplementary material. There, we demonstrate that inclusion of the thermodynamic drive for covalent bond formation does not alter our conclusions.

The studies of short DNA/RNA duplexes report bimolecular association rate constants on the order of ≈ 10^6^ − 10^7^M^−1^sec^−1^ at room temperature and high salt concentration (Na^++^ ≈ 1M) for DNA [70]. Prebiotic monomer concentrations are estimated to have ranged from micromolar to millimolar levels [71]. Based on this, we assume the base pair formation rate p=2 sec^−1^ at a monomer concentration of 2 *µM*. The base pair formation is entropically driven and is therefore taken to be the same for both correct and incorrect pairings. However, the dissociation rate of incorrect base pairs is known to be about two orders of magnitude faster than that of correct base pairs[18]. In our calculation, we vary the effective base pairing drive 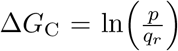, by adjusting the base pair formation rate *p*, keeping the correct base pair dissociation rate *q*_*r*_ = 1 sec^−1^ and the incorrect base pair dissociation rate *q*_*w*_ = 100 sec^−1^ constant [18].

#### ii. Kinetic parameters

Our model requires only one free kinetic parameter, *α*, which quantifies the amount by which a base pair catalyzes the rate of formation/dissociation of a base pair to the 5^*′*^-end of the template. The increase in the kinetic barrier to the 3^*′*^-end of the template is simply assumed to be 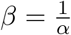, to keep the number of parameters low. The kinetic variable *v*, denoting the rate of transition from the either of the two final states 11111 or 11211 to the covalent bonded states, is expressed in terms of the mean first passage time for the Markov chain to reach the correct final state 11111, starting from the initial state 10000. Thus, we define a dimensionless covalent bonding rate 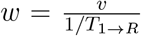, such that *w* ≈ 1 would imply that the covalent-bonding step proceeds at a rate comparable to that of the base pairing rate (see “Methods” above). We scanned *α* and *v* over several orders of magnitude to assess their effect on error correction (Fig.6 in the section 4.II). We show that kinetic discrimination emerges only when *v* begins to become comparable or faster than the base-pairing timescale (i.e. *w* ≈ 1).

**Figure 4:**
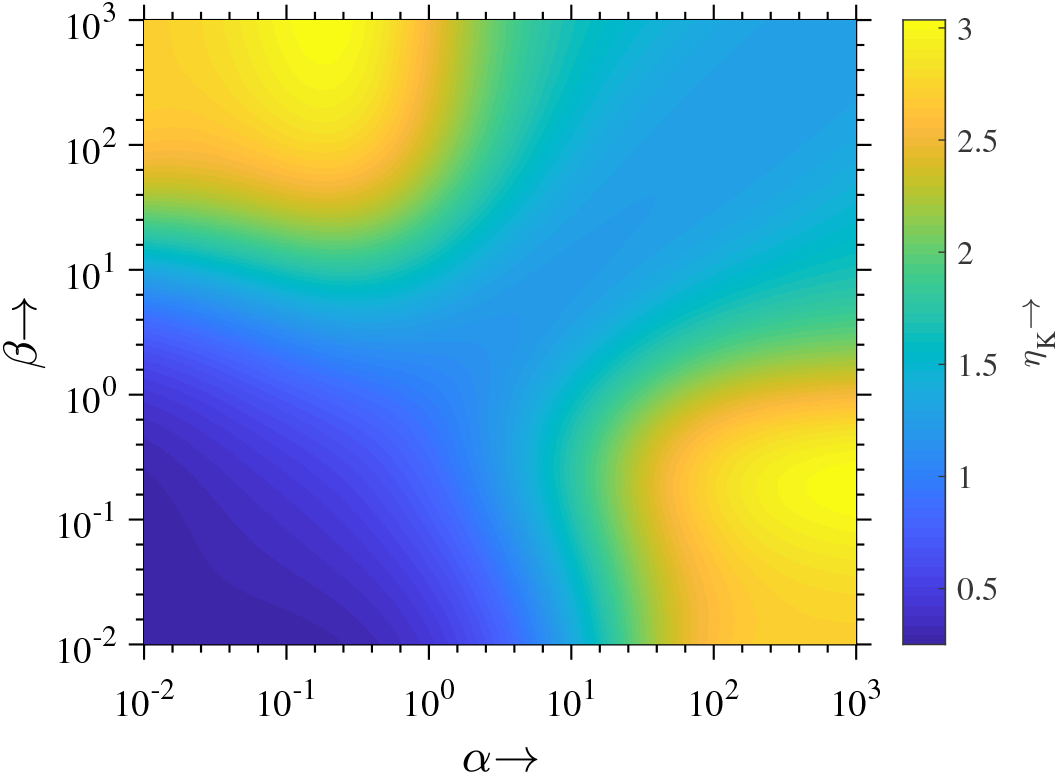
Accuracy ratio (*η*_K_) as a function of the kinetic parameters *α* and *β*, for a linear template comprising *N* = 5 nucleotides and an incorrect base pair at the third location. The base pair formation and dissociation rates are set at 2 sec^−1^ and 1 sec^−1^, respectively. Each point on the plot corresponds to a unique pair of *α* and *β* values, with the color intensity indicating the magnitude of the accuracy ratio. The plot exhibits two distinct maxima, where *α* ≠*β*, positioned symmetrically away from the diagonal. An optimal accuracy ratio is achieved when the cooperative kinetic interactions between the base pairs are asymmetric, whereas symmetric cooperativity along the diagonal (*α* = *β*) results in the lowest possible accuracy ratio. Unidirectional strand construction enables error correction, within this model.

**Figure 5:**
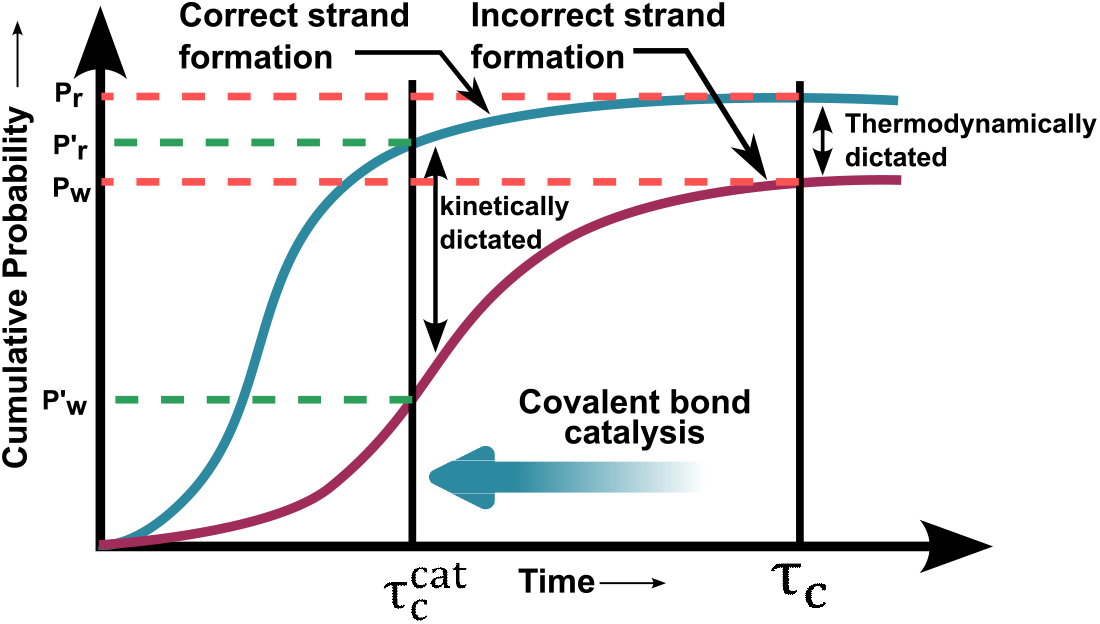
A schematic illustration of how covalent bond catalysis leads to error correction. The diagram also illustrates how a transient time difference between correct and incorrect strand construction is converted into persistent structural order through irreversibility-aided error correction. Two curves, a blue one depicting the cumulative probability of formation of correct strand as a function of time, and a red one depicting the time variation of cumulative probability of incorrect strand formation, are shown. These base pair formation/dissociation events are reversible, and are presumed to happen at rates much faster than the timescale of the figure. Due to asymmetric cooperativity-induced kinetic discrimination between correct and incorrect base pairs, a transient time delay arises between the correct and incorrect strand formation, with incorrect strand formation transiently lagging behind, as illustrated by the slow rise of its cumulative probability curve. When the covalent bond formation between two nucleotides on the daughter strand is not catalyzed, the expected time *τ*_*c*_ for the formation of the covalent bond would be very large, by which time, the above transient time difference would have ceased to exist. This uncatalyzed covalent bond formation time is denoted by a vertical line towards the right. Therefore, uncatalyzed covalent bonds irreversibly stabilize base pairs whose error frequencies, calculated as 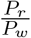, are determined by thermodynamics alone. This is the regime most non-enzymatic DNA replication experiments operate in. When the covalent bond is catalyzed by a polymerase or a metal ion, the expected time for its formation decreases to 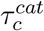, denoted by the left vertical line. If this time is within the time window when the transient kinetic difference between correct and incorrect base pairs persist, the irreversible covalent bond formation can discriminate between the correct and incorrect base pairs better than the thermodynamically-imposed limit, and freezes-in the correct base pair. The accuracy ratio in this regime is given by 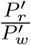.

**Figure 6:**
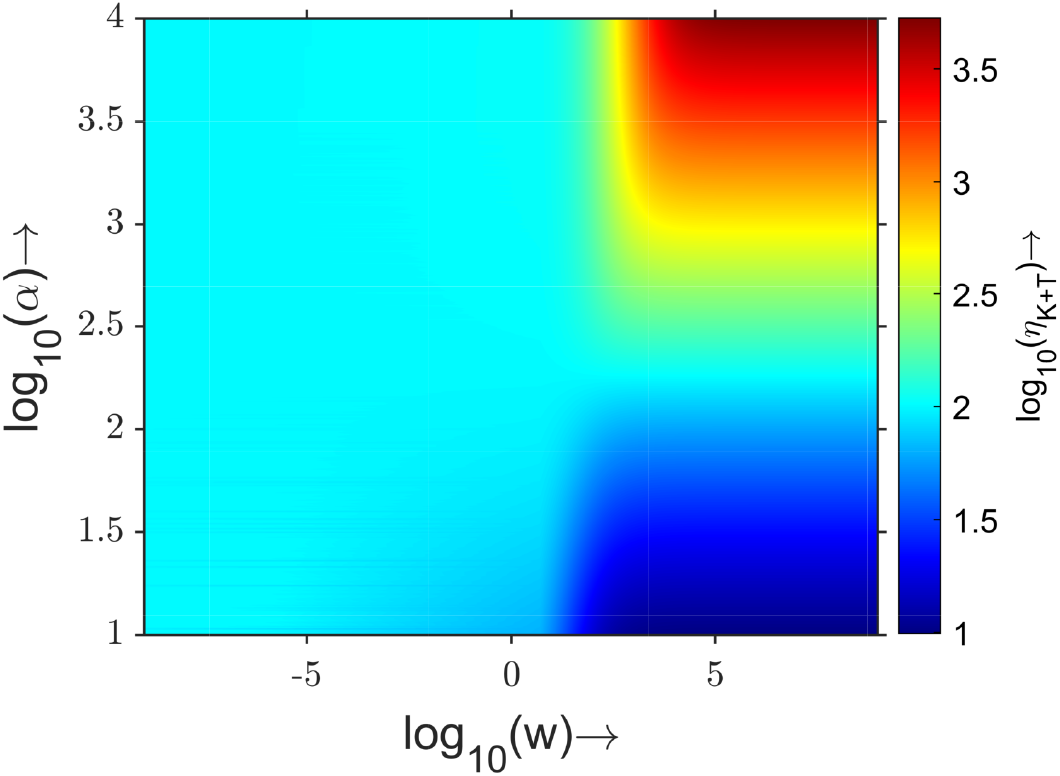
Variation of accuracy ratio (*η*_K+T_) as a function of kinetic asymmetry parameter *α* and dimensionless covalent-bond formation rate *w*, with *β* = 1*/α, p* = 2 sec^−1^, *q*_*r*_ = 1 sec^−1^ and *q*_*w*_ = 100 sec^−1^. The dimensionless covalent-bond formation rate is defined by 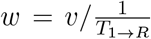, where *v* denotes the covalent-bond formation rate and *T*_1→*R*_ is the conditional mean first-passage time to the fully correct bonded state 11111. When *w <* 1, covalent-bond formation is slower than base-pairing dynamics, and the system exhibits only thermodynamic discrimination, resulting in a thermodynamically dictated accuracy of 10^2^. In this regime, the irreversible covalent bonding is too slow to utilize the time difference between the correct and incorrect daughter strand construction. For *w >>* 1 and large *α*, the irreversible covalent-bond formation step rapidly stabilizes the faster correctly paired state, effectively resolving the time lag between correct and incorrect strand formation, leading to accuracy value exceeding the thermodynamic limit.

To find an experimentally meaningful value of *α*, we utilize the experimentally determined ratio of rates of dissociation of incorrect to correct base pairs. This ratio is determined to be of the order of 10^3^ in [72], for T7 DNA polymerase. Within our model, this ratio is MFPT_11100→11000_*/*MFPT_11200→11000_. In Fig.S5, we show the variation of this ratio as a function of *α*, from which we derive a value of 5 *×* 10^3^ for *α*. In their theoretical examination of kinetic versus thermodynamic contribution to error correction, Sartori *et al* [73] employed the above experimental data to constrain their model and used 3 *×* 10^3^ − 6 *×* 10^4^ (corresponding to a range of 8 - 11 for their kinetic discriminatory factor *δ* between correct and incorrect base pairs) for the above-defined ratio, similar to the value we have used.

The value of alpha used to reproduce the accuracy of base selection process can also be motivated by the experimentally observed fact that the ratio of rates of all successive base pair formations to the rate of initial or the first base pair formation (which is unassisted by its neighboring base pairs) on a template, during the annealing of two single strands of DNA, is of the order of 10^2^ to 10^3^ [74–76]. In other words, the rate of zippering of the two single strands is about 10^3^ times faster than the initial nucleation event. Here, we have assumed that the rate enhancement due to neighborhood *kinetic* influence to be of a similar order.

## 4. Results

This section is organized as follows. First, in part I, we allow the kinetic parameters *α* and *β* to vary *independently* to demonstrate that asymmetric cooperativity, and hence unidirectional strand construction, play an important role for error correction. We include the state 00000 as the initial state only for this demonstration, in order to maintain the left-right symmetry (for the rest of the analysis, we initiate the Markov chain from the 10000 state, and exclude 00000 state from the state space). We show that kinetic discrimination, defined as the presence (absence) of asymmetric cooperativity in correctly (incorrectly) formed base pairs, is most effective in correcting errors when cooperative kinetic interactions between neighboring base pairs are highly asymmetric, and minimal when symmetric (*α* = *β*).

In part II, we introduce covalent bond formation between the base paired nucleotides by incorporating irreversible transition states into the reaction network. We show that catalysis of the covalent bond formation freezes and amplifies the kinetic differences between correct and incorrect base pairing – even if the covalent bond formation step itself is non-discriminatory. Kinetic discrimination vanishes at the steady state if this irreversible transition is removed from the reaction network. In part III, we show that there is a trade-off between strand construction speed and accuracy in our model, an observation recorded in nearly every other error correction scheme proposed and tested thus far. This trade-off implies the existence of an optimal effective base-pairing drive at which the error correction is maximal. We argue that this trade-off may result in a built-in capability for adaptive tuning of error rates in primordial self-replicators, in response to variations in resource availability or environmental conditions, thereby enhancing the rate of evolutionary search for optimal processes and configurations.

### I. Unidirectional strand construction is essential for error correction

In our previous work[43], we argued that unidirectional strand construction arises from asymmetric cooperativity, where a base pair influences the formation and stability of neighboring base pairs unequally on either side, promoting growth in a single direction. We showed that this directional bias resolves conflicting requirements of low kinetic barrier for fast nucleotide incorporation and high kinetic barriers for base pair stability to promote covalent bonding, giving such heteropolymers an evolutionary advantage over symmetric (bidirectional) replicators.

Our contention that asymmetric cooperativity (and unidirectional daughter strand construction as its consequence) leads to error correction can be explained as follows. As shown in Fig.2, a correct base pair at *m*^*th*^ position catalyzes the formation of another base pair towards its right at (*m* + 1)^*th*^ position (template’s 5^*′*^ direction) by lowering the latter’s kinetic barrier, thereby extending the strand construction further. This stabilizes the correct base pair at *m*^*th*^ position, due to inhibition from the (*m* + 1)^*th*^ base pair, resulting in a longer lifetime for the *m*^*th*^ correct base pair. However, an incorrect base pair at *m* has a low probability to extend the daughter strand construction to the right, due to its inability to reduce the kinetic barrier towards its right (5^*′*^-end direction), according to our central premise. Therefore, the kinetic barrier and hence the stability of the incorrect base pair at *m* is lowered, due to the absence of a stabilizing base pair at (*m* + 1) and the presence of a destabilizing base pair at (*m* − 1). This results in higher probability for the incorrect base pair to dissociate from the template, resulting in error correction. Thus, the error correction by kinetic discrimination results from unidirectional strand construction, both originating from asymmetric cooperativity. An identical analysis holds if we switch the direction of strand construction from right to left, instead of left to right, by switching the directionality of asymmetric cooperativity.

The foregoing can be quantified by calculating the accuracy ratio *η*, the ratio of the probability for a fully-formed strand with a correct base pair to the probability for the strand with an incorrect base pair at *steady state* (Eq.4), at the predetermined location *m* = 3, as a function of the asymmetric cooperativity parameters *α* and *β*. These two parameters are treated as independent for this demonstration alone. The above probabilities are calculated as explained in the Methods section. We have used an absorbing Markov chain for these calculations, where, the Markov chain is absorbed once it reaches any of the two final states (see below). We calculate the accuracy ratio resulting solely from kinetic discrimination, where a correct base pair influences the kinetic barriers of its neighbors asymmetrically, while an incorrect base pair does not affect its neighbor’s kinetics. We turn off the thermodynamic discrimination between correct and incorrect base pairs by setting *q*_*r*_ = *q*_*w*_. Fig.4 shows the accuracy ratio, *η*_K_, as a function of *α* and *β*, which are varied independently from 10^−2^ to 10^3^. The subscript *K* in *η*_K_ denotes the fact that we have included kinetic discrimination alone. The values of other parameters for generating the plot are *N* = 5 with the incorrect base pair at the third location (Methods), base pair formation rate *p* = 2 sec^−1^ and the dissociation rate *q*_*r*_ = *q*_*w*_ = 1 sec^−1^ for both the correct and incorrect base pairs.

It is clear from Fig.4 that the accuracy ratio *η*_K_ resulting from kinetic discrimination is large when *α* ≠*β*. There are two equivalent maxima, one where a base pair catalyzes its right neighboring base pair formation/dissociation and inhibits the left base pair’s formation/dissociation, and the other where the left neighbor formation/dissociation is catalyzed, and the right neighbor’s, inhibited. These two maxima suggest that when the symmetry of the kinetic influence on the left and right neighbors is broken, discrimination between correct and incorrect base pairs increases. Specifically, when the kinetic influence of a base pair on the left and right neighbors is equal, the error correction is minimal. This demonstrates the fundamental connection between unidirectional replication and error correction. Apart from the evolutionary advantage of unidirectional replication identified in [43], asymmetric cooperativity is shown here to provide an additional advantage of error correction. Thus, we conclude that unidirectional daughter strand construction constitutes a key mechanistic requirement for achieving error correction within this model.

### II. The role of rapid, irreversible covalent bond formation in error correction

As explained in “Methods” section, we employed two approaches to model the base pairing of nucleotides on a template strand during DNA/RNA self-replication: non-absorbing and absorbing Markov chains. In the case of non-absorbing Markov chain approach, the hydrogen bonds between nucleotides are allowed to break even after the chain reaches the terminal state 11111. This makes the Markov chain reversible. In the non-absorbing case, such reverse transitions are not allowed, and once the chain reaches the final state it cannot return back. The second case thus implements a strong non-equilibrium drive towards daughter strand formation. A detailed comparative analysis of these two approaches is provided in the supplementary material (section V). It is important to note that, while the non-absorbing Markov chain loses its ability to kinetically discriminate between correct and wrong basepairs at steady state, the absorbing Markov chain enables it, due to its inherent non-equilibrium nature. Only in the non-absorbing case does the replication accuracy improves beyond the thermodynamically dictated limit and attain values observed experimentally.

In this article, we identify the source of the above-mentioned non-equilibrium drive that prevents hydrogen bond dissociation as the covalent bond formation between nucleotide neighbors hydrogen bonded to the template. Since covalent bond formation, involving hydrolysis of triphosphate bond, is highly exergonic (ΔG = −7kcal/mol = −11.8*k*_*B*_*T*) [56], this reaction can be treated as irreversible. Once covalent linkages are established, backward transitions (such as pyrophosphorolysis) become effectively impossible and do not contribute to error correction. This assumption is supported by experimental observations, such as those reported by Wong et al.[77], who noted that “the rate of pyrophosphorolysis on a mismatched 3^*′*^-end is undetectable, indicating that pyrophosphorolysis does not play a proofreading role in replication”. How is this irreversibility used for error correction?

As explained earlier in the article, reaching a final state with an error takes longer than reaching the correct final state. This time difference between wrong and correct daughter strand formations can be used for error correction if the faster correct daughter strands are immediately secured through the irreversible covalent bond formation. The rate of covalent bond formation must be high such that it can *resolve* the difference between the correct and wrong strand formation times. To count seconds, one must have a timer that ticks at least once every second, if not faster. The idea that a transient temporal difference can be converted into a persistent structural order, through irreversibility, is schematically illustrated in Fig.5.

To capture this mechanism within our mathematical framework, we introduce two additional terminal states: **R** and **W**, representing covalently bonded correct and incorrect final states, respectively. We make the above two states absorbing, instead of the earlier absorbing states 11111 and 11211. Since we only care about incorporation of the incorrect/correct nucleotide at the *m* = 3 location into the daughter strand, the terminal states **R** and **W** represent daughter strands with a covalent bond between the *m* = 3 and the previous *m* = 2 nucleotides. The covalent bonds between previous nucleotides are assumed to have formed, while the bonds between nucleotides at m=3 & 4 and m=4 & 5 are assumed to have not formed yet, although nucleotides are base paired to the template at these locations. The justification for these assumptions are provided in the Methods section. These terminal states are connected to the fully base paired configurations 11111 and 11211, with the dimensionless forward transition rates given by 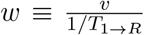, where *T*_1→*R*_ is the conditional mean first-passage time to the correctly base paired covalent bonded state **R**. This implies that the absolute forward transition rate *v* from the non-covalent-bonded to the covalent-bonded states are measured in terms of the conditional mean first passage time to reach the correct state **R** from the initial state. we calculate the accuracy ratio *η*, defined in Eq.4, to quantify the changes in error correction when covalent bonding is added to the model.

In Fig.6 we plot the accuracy ratio *η*_K+T_ resulting from the combined discrimination (kinetic and thermodynamic) as a function of the kinetic asymmetry parameter *α* and the dimensionless covalent bond formation rate *w* (with *β* = 1*/α*). Two distinct regimes emerge: When *w <<* 1, covalent bond formation rate *v* is much slower compared to the rate of daughter strand construction with all correct base pairing (11111), 1*/T*_1→*R*_. Covalent bond formation is simply not fast enough to discriminate the time difference between the correct and incorrect daughter strand formations, in this regime. Here, *η*_K+T_ remains ∼ 10^2^ for all *α*, the maximum discrimination possible from thermodynamics of correct/incorrect base pairing alone. As *w* → 1, kinetic effects on the error correction begins to appear and *η*_K+T_ rises above the thermodynamic limit with *w* and *α*. Covalent bond catalysis (*w*) and unidirectionality of strand construction (*α*) both contribute to the kinetic discrimination: unidirectionality ensures maximal time lag between correct and incorrect daughter strands, whereas, higher covalent bond formation rate is now capable of resolving this time lag. This leads to kinetics-enabled enhancement in error correction with increasing *w* and *α*, with *η*_K+T_ reaching ∼ 4 *×* 10^3^.

To further clarify the mechanistic origin of this behavior – and to explicitly differentiate the contributions of thermodynamic and kinetic discrimination, the subsequent analysis examines how the accuracy ratio varies with the covalent bond formation rate *w* under three distinct regimes: the kinetic discrimination regime, the thermodynamic discrimination regime, and the combined regime. Fig.7 shows the effect of covalent bond catalysis *w* on the accuracy ratio *η*. For this and all other analyses henceforth, we choose *α* = 5 *×* 10^3^, a value that we derived by matching the dissociation rate ratio between correct and incorrect base pairs from experiment [72] to our model (Methods).

**Figure 7:**
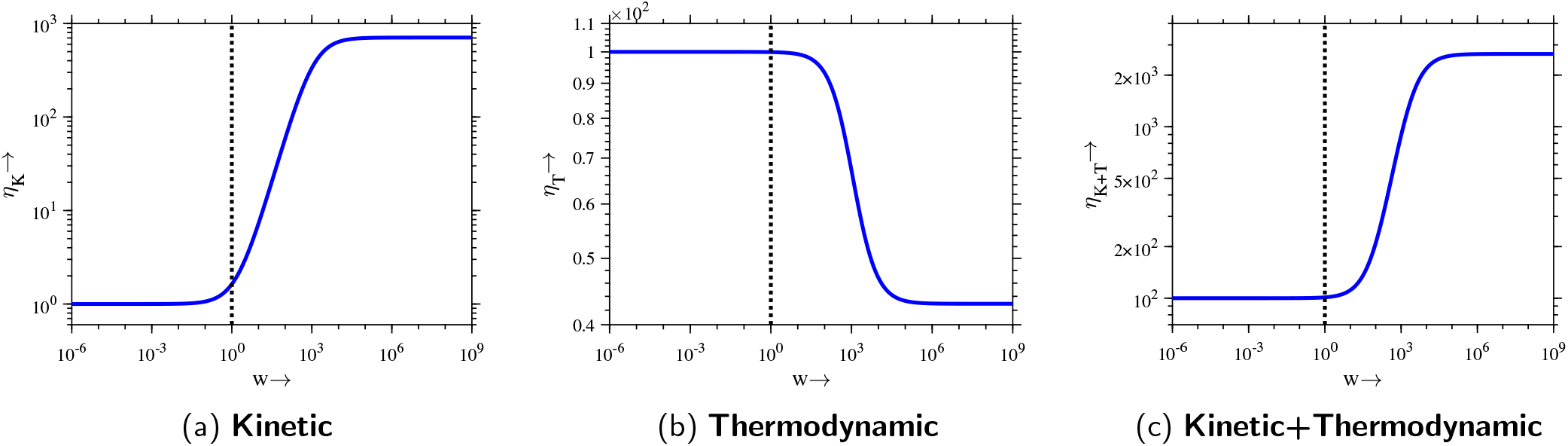
Illustration of accuracy ratio *η* as a function of the covalent bond formation rate *w* under three discrimination scenarios. In each panel, the vertical dashed line marks 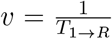: (a) Kinetic discrimination (*q*_*r*_ = *q*_*w*_ = 1 sec^−1^, *α* = 5 *×* 10^3^ only for correct base pairs): At very low rates of covalent bond formation, i.e.,(*w <<* 1 or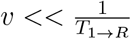, the slow covalent bond formation event is unable to resolve the comparatively much smaller difference between the base pairing timescales of the correct and incorrect nucleotides. This leads to *η*_*K*_ ≈ 1. When the rate of covalent bonding, *v*, matches or exceeds the rates of base pair formation and dissociation,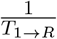, the covalent bonding event begins to become sensitive to the time difference between the correct and incorrect base pairing dynamics. This location, *w* = 1, is highlighted by a dashed line. From this point on, the covalent bond begins to preferentially incorporate the correct nucleotide over the incorrect one by locking-in the faster correct nucleotide, error correction rapidly improves, and saturates at a value dictated by the kinetic parameter *α*. (b) Thermodynamic discrimination (*q*_*r*_ = 1 sec^−1^, *q*_*w*_ = 100 sec^−1^, *α* = 5 × 10^3^ for both correct and incorrect base pairs): Increasing the covalent bonding rate *w* in this regime leads to reduced accuracy. When bond formation rate is slow (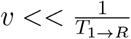), the system has sufficient time to explore multiple configurations and settle into a thermodynamically favored state, leading to a discrimination factor (*q*_*w*_*/q*_*r*_ = 100) determined entirely by thermodynamics. When the irreversible covalent bond formation rates increase beyond 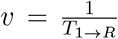, the time for exploration of multiple configurations is minimized, the now-nondiscriminating kinetic factors dominate the time evolution of the Markov chain, leading to cessation of error correction. (c) Combined discrimination (*q*_*r*_ = 1 sec^−1^, *q*_*w*_ = 100 sec^−1^, *α* = 5 *×* 10^3^ only for correct base pairs): When both kinetic and thermodynamic mechanisms are at play, the behavior of the system mirrors that observed in the kinetic regime with high accuracy ratio *η*_K+T_. This highlights the dominant role of kinetics in error correction and suggests that rapid and irreversible covalent bond formation ensures high fidelity.

In Fig.7, the accuracy ratio is depicted as a function of the covalent bond formation rate *w*, across these three distinct regimes. The sharp distinction between the kinetic and thermodynamic regimes can be seen in the Figs.7a and 7b: when the covalent bonding rate increases, kinetic discrimination helps in increasing the accuracy ratio, whereas, thermodynamic discrimination hinders the accurate self-replication. When the covalent bonding rate is high, discriminating the time difference between the early correct sequence and the later-arriving wrong sequence becomes more effective. Therefore, higher covalent bonding rates lead to better error correction. This can be seen from the knee point in Fig.7a, where the accuracy ratio begins to increase, at 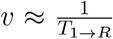. As soon as *v* exceeds 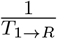, the covalent step irreversibly “locks-in” the faster-arriving Markov chain corresponding to the correct daughter strand, so any kinetic advantage amplifies into a large accuracy gain.

On the other hand, the behavior of *η* in Fig.7b can be understood by considering the role of *w* as the rate controlling the transition from intermediate states to the absorbing states (**R** and **W**). When *w* is small, the absorption process occurs slowly, allowing the system to dwell longer in the transient states. This extended residence time facilitates the establishment of a quasi-equilibrium among the intermediate states, such that the occupation probabilities reflect thermodynamic biases. This regime is nearly equivalent to the non-absorbing Markov chain approach mentioned earlier. In contrast, when *w* is large, the system transitions rapidly to the absorbing states, significantly reducing the time available for equilibration among the transient states. As a result, the accuracy ratio is increasingly determined by the kinetic pathways rather than equilibrium distributions, which are set to be non-discriminatory in this regime. In the combined regime, Fig.7c, which includes both thermodynamic and kinetic discrimination, the behavior of *η* is similar to that of the kinetic regime, pointing to the importance of kinetics in error correction. We provide a considerably simpler, analytically tractable model of daughter strand construction in the supplementary material (Section. II), where, we mathematically demonstrate the lack of error correction for very small values of *w*, the relative covalent bonding rate, and show that the error correction approaches values observed in passive polymerases for high values of *w*.

With the above demonstration, we conclude that covalent bond catalysis is a crucial factor in accurate construction of daughter strands. This mechanism alone can account for the ability of thermodynamically passive polymerases to correct replication errors without any external energy input, thereby potentially resolving the paradox highlighted in the introduction. Furthermore, it also accounts for the high error rates observed during non-enzymatic DNA self-replication. With a thermodynamic discrimination between the correct and wrong base pairing of 10^−2^, the maximum error correction achieved in our model is about ∼ 1*/*(3.9 *×* 10^3^) = 2.56 *×* 10^−4^, which corresponds with the experimentally observed error ratio 10^−4^ when employing thermodynamically passive polymerases [17, 23]. In this context, passive polymerases function primarily to catalyze covalent bond formation, a process that facilitates error correction as explained above. The energy supply for passive base selection comes from the irreversible covalent bond formation, within our model. The above conclusions are not specific to templates of a certain length. In Supplementary material (Section.VI), we extend the analysis to templates of lengths N = 6, 7 and 8 while keeping the error site fixed, and show that it yields qualitatively similar trends.

In the primordial scenario, when polymerases were not available, how was the accuracy of daughter strand construction maintained? The rate of covalent bond formation in the primordial scenario was possibly very slow, of the order of minutes to hours, whereas, the rate of base pairing was of the order of seconds [78]. These values suggest that a catalyst for covalent bond formation must have been present in the primordial scenario to catalyze daughter strand polymerization and thus maintain its accuracy. Metal ions (primarily *Mg*^++^), which are present in all extant DNA polymerases without exception [79] and perform crucial error correction functions, might have assisted covalent bond formation in DNA/RNA or their precursors in the primordial scenario, most probably through electrostatic interactions [79–81]. The foregoing implies that a search for efficient inorganic prebiotic catalysts for polymerization of DNA/RNA precursors may also help solve the replication accuracy problem in the primordial scenario.

### III. Thermodynamic regulation of the speed–accuracy trade-off in strand construction

The general observation of an inverse relationship between product formation rate and product accuracy during error correction is pervasive in the literature [12, 73, 82–85]. In the paradigmatic Hopfield-Ninio scheme, as the product formation rate (*w* in [3]) decreases, the error correction improves. Variations on this basic scheme show interesting behavior, such as the observation of the presence of an optimal product formation rate at which the error is minimized[48–51]. Our model exhibits a similar non-monotonic behavior when the base-pair formation rate *p* is varied with fixed *q*_*r*_ = 1 sec^−1^ and *w* = 10^5^, thereby tuning the effective base-pairing drive Δ*G*_c_, which quantifies the free-energy bias associated with correct base-pair association and dissociation.

This effective base-pairing drive determines the balance between two competing processes that are influenced by *p*: the *stabilization* of the incorrect base pair at *m* by the newly added neighboring base pair at (*m* + 1), on the 5^*′*^ side, and the incorrect base pair’s *dissociation*, catalyzed by a correct base pair at (*m* − 1), on the 3^*′*^ side. The outcome of this competition gives rise to an inherent speed-accuracy trade-off in the model upon inclusion of kinetic discrimination, where high base-pair formation rates favor rapid extension but increase the likelihood of retaining errors, while very low bond formation rates decrease the lifetime of base pairs, which reduces their ability to reduce the kinetic barrier of the incorrect base pair towards their 5^*′*^-end, leading to reduced error correction. An optimum lies between these two limits, where the drive is just enough to provide long enough lifetime for the *m* − 1 neighbor of the incorrect base pair at *m*, enhancing error correction, but is not large enough to allow the uncatalyzed formation of a stabilizing base pair neighbor of an incorrect base pair at the (*m* + 1) location. This trade-off is quantitatively illustrated in Figs.8a and 8c, which shows the accuracy ratio *η*_K_ as a function of the thermodynamic drive 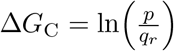.

**Figure 8:**
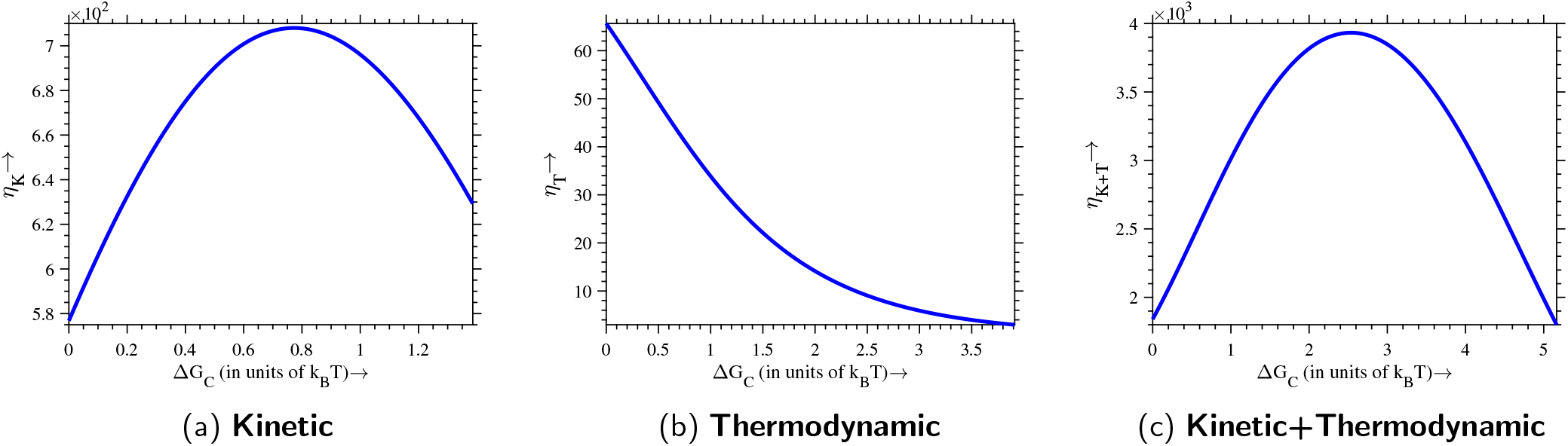
Accuracy ratio *η* as a function effective base pairing drive Δ*G*_C_. Panels (a) and (b) correspond to purely kinetic (*q*_*r*_ = *q*_*w*_ = 1 sec^−1^, *α* = 5 × 10^3^ only for correct base pairs) and purely thermodynamic (*q*_*r*_ = 1 sec^−1^, *q*_*w*_ = 100 sec^−1^, *α* = 5 × 10^3^ for both correct and incorrect base pairs) discrimination, respectively, while panel (c) shows the combined regime incorporating both kinetic and thermodynamic discrimination (*q*_*r*_ = 1 sec^−1^, *q*_*w*_ = 100 sec^−1^, *α* = 5 *×* 10^3^ only for correct base pairs). Below an optimal effective base-pairing drive corresponding to the peak, the base pair formation is low, and the catalytic influence of the (*m* − 1) neighbor of the incorrect base pair at *m* is lower, leading to lower error correction. On the other hand, above the optimal effective base-pairing drive, the formation of correct base pairs next to incorrect base pairs occurs rapidly, stabilizing the incorrect base pairs and increasing the error probability. An optimal error correction lies between these two regimes.

Fig.8a illustrates the dependence of *η* on the effective base pairing drive Δ*G*_c_ in the case of purely kinetic discrimination (*q*_*r*_ = *q*_*w*_). It can be observed that the accuracy exceeds unity even at Δ*G*_c_ = 0. We attribute this to the non-equilibrium condition imposed by the absorbing Markov chain approach. Within this approach, the presence of an absorbing final state introduces a unidirectional bias in the dynamics, effectively enforcing irreversibility in the daughter strand construction process. Consequently, the system inherently disrupts detailed balance and maintains a nonzero net flux toward the final (absorbing) state, even in the absence of an explicit thermodynamic drive. As a result, the system remains out of equilibrium and supports a finite degree of error correction even at Δ*G*_c_ = 0. In Fig.8b, *η*_T_ exhibits a monotonic decrease with increasing Δ*G*_C_. This indicates that in the absence of kinetic assistance, increasing Δ*G*_C_ accelerates the incorporation of both correct and incorrect nucleotides indiscriminately, thereby reducing the opportunity for error correction. Fig.8c illustrates the accuracy ratio *η*_K+T_ under conditions of combined discrimination, where both kinetic and thermodynamic discrimination contribute to error correction. The explanation for the behavior of *η*_K+T_ is given above. Overall accuracy is enhanced compared to either regime independently, indicating that the two mechanisms synergistically enhance fidelity.

These observations highlight the distinct but reinforcing effects of kinetic and thermodynamic discrimination in enhancing the accuracy of daughter strand construction. Kinetic discrimination operates by introducing temporal delays at error-prone steps, thereby allowing incorrect incorporation to be selectively rejected before finalization. Thermodynamic discrimination, by contrast, relies on energetic differentials-quantified as 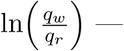 to thermodynamically favor correct base pairings over incorrect ones. The interplay between these mechanisms gives rise to a finely tunable accuracy landscape governed by the effective base pairing drive 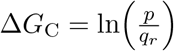.

Finally, because Δ*G*_c_ captures only the free-energy bias associated with hydrogen-bond formation and dissociation of correctly paired nucleotides, it does not constitute the total thermodynamic drive underlying daughter-strand construction, which includes the irreversible covalent bond formation. Therefore for completeness, we also analyzed the behavior of *η* with respect to the total thermodynamic drive in terms of total entropy production rate, which allows us to assess how accuracy ratio *η* responds not only to the correct base-pairing bias but to the full thermodynamic drive acting on the system. The results, presented in the supplementary material (Section. III), shows a drive–accuracy trade-off very similar to the one shown above: as the base-pair formation rate *p* increases, the total entropy production rate concomitantly rises, leading to an initial enhancement in the accuracy *η*_K+T_ up to an optimal value, beyond which further increase in total entropy production rate results in low accuracy.

## 5. Experimental support

Our model captures several key phenomena observed in experimental studies of error correction in template-directed polymerization. In the following, we discuss experimental and theoretical findings in the literature that support and validate the assumptions underlying our model.

### I. Stalling

Stalling denotes the slow-down of the construction of daughter strands with an incorrect base pair, compared to the rate of construction of an error-free daughter strand. In our approach, stalling is demonstrated by evaluating the conditional mean first passage time required to reach the final covalent bonded states – either the fully correct covalent bonded state **R** or the state containing an error **W** – starting from the initial configuration (10000). Our results show that the conditional mean first passage time for reaching the fully correct final state is shorter than that for reaching the error-containing state, as the incorrect base pair inactivates the asymmetric cooperativity, making it less favorable to add new nucleotides to the growing daughter strand. Consequently, the rate of daughter strand construction decreases. This reduction in the extension rate following incorrect base pairing has been well-documented by Rajamani et al. in their study[45]. They use 3^*′*^-amino-2^*′*^, 3^*′*^-dideoxynucleotide-terminated primers to investigate the dynamics of nucleotide incorporation during *non-enzymatic* polymerization, particularly looking at what happens when there are incorrect base pairs (mismatches) at the primer-template junction. Their findings indicate that the polymerization process halts in the presence of incorrect bases, allowing accurate replication to proceed more efficiently than erroneous replication. Further support for this reduction in extension rate due to incorrect base pairing comes from the work of Leu et al.[46]. They use gel electrophoresis and mass spectrometry to quantitatively measure extension rates and mutation profiles in *non-enzymatic* nucleic acid strand synthesis. Their results demonstrate that the rate of daughter strand construction in the presence of incorrect (mismatched) base pairs is 10 to 100 times slower compared to error-free conditions. Göppel et al.[47] offer further evidence for stalling induced by incorrect base pairing through their investigation of *enzyme-free* template-directed polymerization dynamics via kinetic modeling. In their study, they introduced mismatches at the ligation site to assess the impact of non-complementary nucleotides on the polymerization process. Their findings indicate that the presence of mismatched base pairs results in a marked deceleration of strand extension, a phenomenon they termed kinetic stalling.

In Fig.9, we present the stalling quotient, defined as 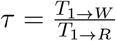 obtained under purely kinetic discrimination as well as in combination with thermodynamic discrimination, plotted as a function of the effective base pairing drive ΔG_*C*_. We observe that in both the plots, *τ* exhibits a non-monotonic dependence on the effective base pairing drive ΔG_*C*_, similar to the behavior of *η* in Figs.8a and 8c: it increases initially, reaches a maximum, and then decreases. Importantly, across the entire range of Δ*G*_*C*_, the stall quotient *τ* remains greater than one, indicating that the conditional mean first passage time to the incorrect final state consistently exceeds that of the correct final state. Furthermore, the peak position of the stall quotient *τ* aligns closely with the optimum observed in the accuracy plots shown in Figs.8a and 8c. This correspondence supports the proposed mechanistic link between stalling induced by incorrect base pairing and the increased likelihood of subsequent dissociation of the incorrect base-pair, thereby enhancing the accuracy of the daughter strand construction, quantified by the accuracy ratio (*η*).

**Figure 9:**
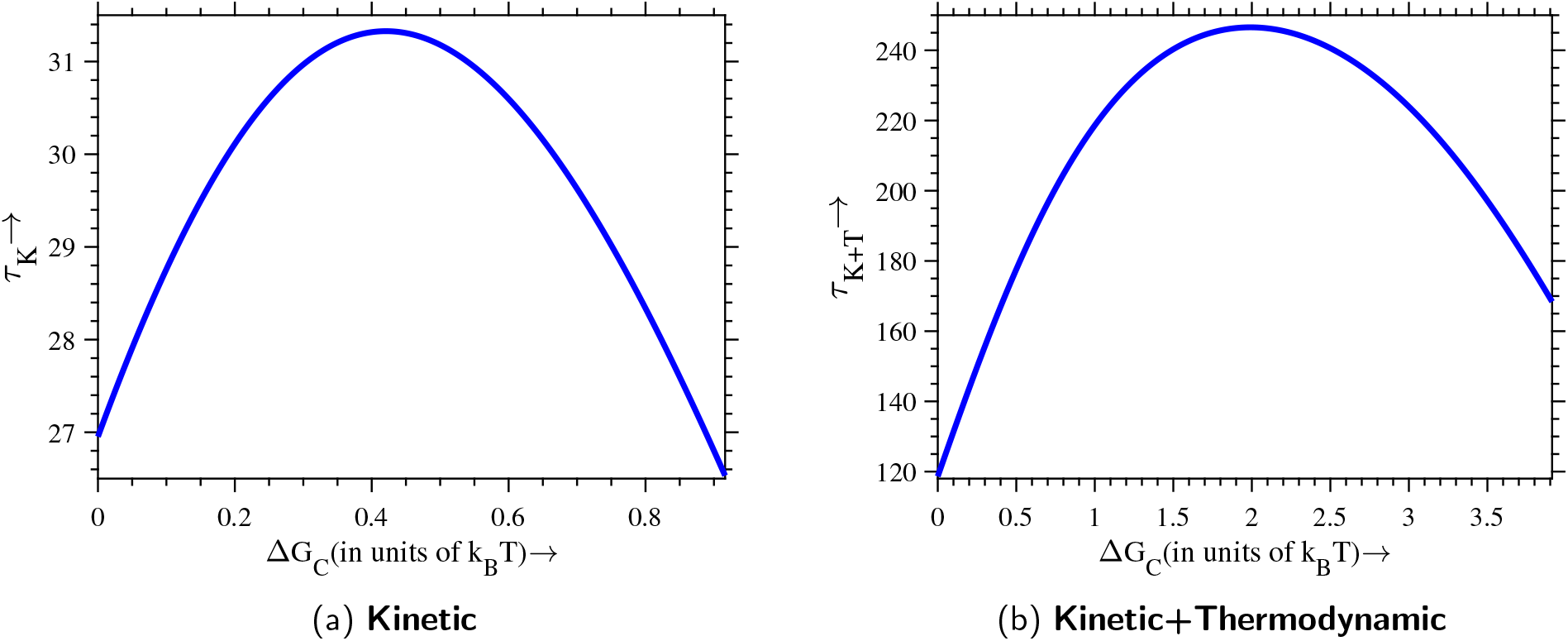
Stalling quotient 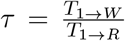 as a function of effective base pairing drive Δ*G*_*C*_. The parameters used to generate these plots are the same as that of figs.8a and 8c. The plots show the conditional mean first passage time to the error-containing final state (**W**) relative to that for the error-free final state (**R**), under (a) purely kinetic discrimination and (b) combined kinetic and thermodynamic discrimination. In both cases, *τ* exhibits a non-monotonic dependence on Δ*G*_*C*_, peaking at intermediate values. The quotient *τ* remains greater than one throughout, indicating that incorrect base pairing leads to a slowdown in daughter strand construction at all drives. This maximum closely corresponds to the optimal accuracy points in figs.8a and 8c, supporting a mechanistic link between stalling and improved accuracy.

### II. Fraying

When the formation of the new base pair is stalled due to an incorrect base pair at *m*, both the (*m* − 1)^*th*^ and *m*^*th*^ base pairs in Fig.2 at the growing front remain unstable and are prone to dissociate, due to the low kinetic barriers at their respective locations. Dissociation of any of these two base pairs can precipitate a sequential dissociation of other base pairs in the 3^*′*^ direction, leading to strand *fraying*. Therefore, the presence of an incorrect base pair increases the likelihood of fraying of the daughter strand from the template strand, in our model. This phenomenon is experimentally observed in [52], where Fernandez-Leiro et al. used cryo-electron microscopy (cryo-EM) and nuclear magnetic resonance (NMR) spectroscopy to study the structural dynamics of a mismatched DNA substrate during high-fidelity replication. They found that the presence of an incorrect base pair increases the fraying of the DNA duplex by one additional base pair at the 3^*′*^ terminus of the daughter strand, leading to a distorted conformation that facilitates the transfer of the DNA strand to the exonuclease active site for proofreading. Their findings indicate that mismatch-induced fraying acts as a self-correcting mechanism, where the DNA itself enhances its error correction by increasing the instability at the growth front.

Fig.10 illustrates that, in our model as well, the presence of a single mismatch increases the tendency of the strand to undergo fraying. The plots illustrate the probability *P*_10000_(*t*) of reaching the initial one-base-pair state “10000” over time, starting from either the all-correct intermediate state 11100 or the mismatched state 11200. The probability for the state with an error 11200 to revert back to the “10000” state is greater than that of the correct state 11100.

**Figure 10:**
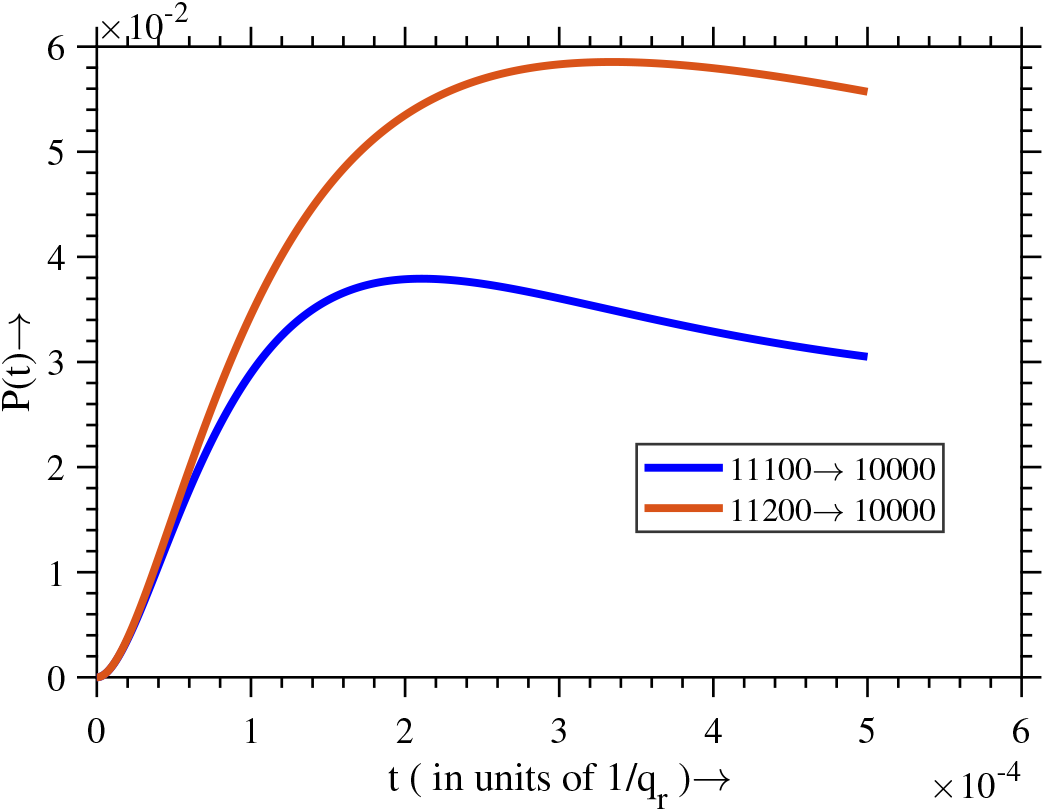
Time evolution of the occupation probability of the state 10000 from two initial conditions: 11100 (all three base pairs 1 − 3 correct) and 11200 (base pairs 1 − 2 correct, position 3 is incorrect). State 10000 represents a frayed configuration. The plot compares *P*_11100→10000_ and *P*_11200→10000_. The probability of reaching 10000 starting from 11200 is larger than from 11100. This disparity indicates that the stand frays much more readily when a mismatch present. The parameters are the same as used for fig.9.

### III. Next nucleotide effects

Upon the formation of a correct base pair at (*m* + 1), next to the incorrect base pair at *m*, in the 5^*′*^ direction of the template, the incorrect base pair is kinetically stabilized, and the extension rate of daughter strand synthesis normalizes, within our model. This stabilization and recovery of the extension rate are due to the asymmetric cooperativity of the newly formed correct base pair, which inhibits the dissociation of the incorrect base pair and facilitates the formation of subsequent base pairs in the 5^*′*^ direction. The ability to recover extension speed following the incorporation of a correct base pair next to an incorrect one (in the 5^*′*^ direction) represents a key feature of our error correction model. This behavior is consistent with the findings of Leu et al.[46], who demonstrated that the addition of a correct nucleotide next to a mismatched base pair can restore the daughter strand extension rate to near-normal levels, indicating a localized stabilization effect that supports continued daughter strand construction.

More over, a correctly paired base immediately downstream of a mismatch tends to lock in the error and prevent its correction. This effect has been characterized in DNA replication: increasing the concentration of the “next correct” nucleotide dramatically reduces proofreading. In classic experiments with *E*.*coli* DNA pol I, Kukel et al. showed that raising the level of the next complementary dNTP increased misincorporation roughly 25-fold at a specific site[53]. They attributed this mutagenic effect to a decrease in exonucleolytic excision: once a correct base is paired after a mismatched end, Pol I’s 3^*′*^ → 5^*′*^ exonuclease is far less likely to remove the mismatch. In other words, the next nucleotide (dATP in their example) stabilized the nascent helix and suppressed proofreading, causing the error to persist.

In contrast, when an incorrect base pair forms next to an existing incorrect base pair in the 5^*′*^ direction, the lack of asymmetric cooperativity associated with the preceding incorrect base pair does not lower the kinetic barrier for the subsequent new base pair in the 5^*′*^ direction. Consequently, the newly added incorrect base pair is not destabilized by the local asymmetric kinetic influence of the preceding incorrect base pair. As a result, the dissociation of the newly formed incorrect base pair becomes less probable, allowing both incorrect base pairs to persist. Thus according to our model, this scenario creates a favorable environment for continued erroneous incorporation, ultimately leading to a cascade of sequential errors. This phenomenon has also been observed experimentally by Leu et al.[46], who reported that an initial mismatch is highly likely to be followed by another mismatch 54 − 75% of the time, resulting in clusters of mismatches and significant kinetic stalling.

### IV. Speed-Accuracy trade-off

Numerous models [12, 73, 82–85] have quantitatively captured speed-accuracy trade-off, where increased accuracy typically comes at the cost of reduced speed. However, our model adds nuance to this conventional relationship. In our model, maximum accuracy is found not at the slowest product formation speed, but at an intermediate speed, determined by the kinetic parameters (Fig.8c). The positive influence of daughter strand construction speed on accuracy below optimal thermodynamic drive (to the left of the peak in Fig.8c) is due to asymmetrically cooperative kinetics: in this regime, daughter strand construction is slow and dominated by sequential nucleotide addition at the growth front, due to catalytic influence of the most-recently formed base pair. The likelihood of forming the (*m* + 1)^th^ base pair is low when the *m*^th^ base is incorrect, due to absent catalytic assistance (Fig.2). This extends the time window for the (*m* − 1)^th^ neighbor to destabilize and dissociate the incorrect base. Increasing the drive parameter *p* in this regime enhances error correction by increasing the (*m* − 1)^th^ base pair formation probability and stability, which increases *m*^th^ incorrect base pair removal efficiency. However, beyond optimal effective base pairing drive (to the right of the peak in Fig.8c), accuracy declines due to incorrect base pair stabilization by the rapidly forming *m* + 1 neighbor, reducing kinetic discrimination advantage. This non-monotonic accuracy profile is consistent with the observations in [48–51].

Ravasio et al.[48] observed this optimality in their minimal non-equilibrium model for molecular order. They simulated Darwinian selection of fast replicators and found that replication speed alone, even in the absence of explicit selection for fidelity, can spontaneously drive the emergence of error-correcting pathways. The evolved networks not only replicated faster but also became more accurate, up to a point — beyond which fidelity declined. This emergent Pareto front separating speed and fidelity parallels the trade-off observed in our results.

Banerjee et al.[49] also observed this non-monotonic speed-accuracy trade-off in their proofreading model using a first-passage time framework. They demonstrated that both speed and fidelity are governed by the enzyme’s internal kinetic parameters. Their analysis reveals a characteristic fidelity-speed curve, where fidelity increases with speed up to an optimum, beyond which error rates rise sharply. Remarkably, the optimal fidelity of polymerases do not coincide with the fastest possible replication; instead, polymerases operate in a biologically tuned regime where speed is favored over maximal accuracy. Their work supports our model’s prediction of a fidelity maximum at intermediate speeds.

A parallel conclusion is presented by Poulton et al.[50], who studied persistent polymer copying in a minimal nonequilibrium framework. They showed that copy–template mutual information — a proxy for replication fidelity — exhibits one or more local maxima at intermediate values of chemical driving (see their Fig.4). At higher driving, fidelity declines due to the loss of kinetic discrimination: when polymerization becomes nearly irreversible, incorrect bases are incorporated as readily as correct ones, diminishing error suppression.

More corroborative evidence emerge from theoretical models of proofreading and enzymatic error correction. Galstyan et al. introduced a spatial proofreading model where diffusion-mediated delays enhance fidelity[51]. By tuning the rate of the driving motor (piston compressions), they found a “resonance” phenomenon: fidelity peaks at a finite driving rate, with lower fidelity at both slower and faster drives (Fig.6D in the paper). Thus the allosteric proofreading engine has an optimal operating frequency for maximum accuracy.

Together, these results support the generality of a speed–accuracy trade-off in polymerization systems. They reinforce the prediction from our model that accuracy is maximized not at the lowest or highest speeds, but at a finely tuned intermediate value, shaped by the interplay between stalling-induced dissociation and catalysis-driven extension.

### V. Adaptive tuning of error rates

Since the base pair formation rate *p* is affected by environmental factors such as temperature, ionic strength, and pH, the optimal drive for error correction—and consequently, daughter strand accuracy, can be modulated by varying the environment(Fig.8c). Such adaptive tuning of error rates would have been particularly advantageous in primordial scenarios, where spatiotemporal fluctuations in thermal, chemical, and other environmental variables were likely more pronounced than they are today[86]. Any non-enzymatic self-replicator capable of adaptively tuning its error rates would have had a significant evolutionary advantage over its competitors in these unpredictable environments. The ability of extant organisms to vary the timing and genomic locations of sequence variability is reviewed extensively in [87]. The preferential localization of essential genes in the mutationally cold regions and that of communication genes that need to respond to environmental variability in the mutationally hot regions in human genome point to the existence of strategies of locally adaptive variations of the mutation rates [88]. Hypermutative phenotypes have been shown to evolve independently in response to ethanol stress in E. Coli[54]. Hypermutative phenotypes of the clinically derived pathogen *A. Baumannii* repeatedly evolved in response to antibiotic treatment, leading to antibiotic resistance [55]. These and many other examples in the literature suggest that organisms are capable of adaptively tuning their error rates, that predictably vary across genomic locations, in response to environmentally induced stress. Our model suggests one possible way such adaptive tuning can emerge: when the environmental stress affects the effective base pairing drive for strand construction, moving it away from its optimal value, the balance between speed and accuracy gets perturbed (Fig.8c), leading to higher mutation rates. The adaptive localization of hypermutative sequences, such as inverted repeats, and within certain genomic locations, such as promoters[89, 90], cannot be modeled within the approach above. However, the model can be extended to include sequence dependence of error rates [91], which should be able to address such adaptive localization of hypervariability.

Collectively, these studies provide evidence supporting our theoretical model and validate our proposed error correction mechanism whereby a reduction in extension rate following incorrect base pairing allows for effective error correction in self-replicators.

## 6. Discussion

A vexing problem concerning the possibility of autonomous emergence of early life has been our inability to demonstrate error-free, non-enzymatic self-replication, and hence faithful transfer of information across generations, under plausible primordial scenarios [92]. A more immediate concern is the lack of clear understanding of the energetics of error correction during DNA replication, specifically of base selection, which does not involve utilization of extraneous energy sources, making the Hopfield-Ninio model inapplicable [17–22]. Base selection is a major contributor to the accuracy of DNA replication, improving the thermodynamically set error rates of about 10^−2^ per base pair to about 10^−4^ per base pair[93]. Other questions include the role of covalent bond formation and of polymerases in error correction[23].

The model we introduced above clarifies the contributions of strand construction kinetics and thermodynamics to error correction, and demonstrates the possibility of non-enzymatic error correction through kinetic discrimination between correct and incorrect base pairs and covalent bond catalysis. We thus identify the source of energy for base selection process as the energy released irreversibly during covalent bond formation itself. No extraneous energy sources are needed for error correction, in contrast to existing models of error correction. This resolves, at least theoretically, the problem of identifying molecules that were capable of discriminating between correct and incorrect base pairs in the primordial scenario, using energy supply. Our answer to this problem is that we do not need a structure-based discriminator[94]: all we need to identify are molecule(s) capable of catalyzing the covalent bond formation in the primordial scenario. Error correction follows, since the asymmetric kinetics already discriminates between correct and incorrect base pairs through time delays in strand construction. This idea considerably simplifies our search for primordial scenarios for evolution of self-replicating informational heteropolymers. With reasonable kinetic and thermodynamic parameters, we reproduce the accuracy of passive polymerases[17] in the base selection process.

Our identification of the source of energy for error correction as the thermodynamic drive for templated strand construction opens up the possibility of modulating the error rates by varying the thermodynamic drive, specifically by varying monomer concentration. Although other environmental parameters, such as temperature, pH and salinity influence the thermodynamic drive, the possibility of living systems tuning the monomer supply rates to optimize the error rates appears more probable [5–7]. This might explain the tight regulation of nucleotide supply rates in extant life forms[95].

Existing theoretical models of error correction in the literature[32–34], apart from utilizing extraneous energy sources, also include multiple parameters to describe the error correction process, thereby allowing degrees of freedom in the model that may not correspond to the ground truth *in vivo*. This leads to the difficult task of explaining how these parameters are collectively fine-tuned by evolution to achieve the incredible levels of accuracy seen in DNA replication. The model we introduced above includes a *single free parameter, α*, describing the asymmetrically cooperative kinetics of strand construction. The remaining are thermodynamic variables, the free energies of correct and incorrect base pair formation and dissociation, whose values and their environmental dependence have been previously determined from extensive studies. This reduction in the parameter space allows for an intuitive understanding of the error correction process without constraining ourselves into specifying regions of parameter space where such an understanding is appropriate. Thus the model provides a simplified yet physically grounded model on error correction mechanism in DNA replication.

One of the fundamental questions regarding origin of life concerns the ability of living systems to create order from non-equilibrium[96]. This order is persistent across time, exemplified by our ability to sequence DNA of long-extinct species, and from the persistence of seeds and spores through inhospitable environments, without any energy supply to maintain order. We contrast this persistence with the ephemerality of the order from non-equilibrium in systems such as convection cells and vortices in turbulent fluids, put forward by the Prigogine’s Brussels School as precursors of the order seen in life. P. W. Anderson’s article[97] clarifies why the examples provided by the Brussels school may be inappropriate for explaining the order found in living systems, none of which show the property of temporal persistence. Here, we provide a model system capable of creating temporally persistent order from non-equilibrium, without requiring the presence of a discriminatory macromolecule, an enzyme. The irreversibility of covalent bond formation, engendered by non-equilibrium created by high initial concentrations of free nucleotide triphosphates, leads to temporal persistence, in our model. This irreversibility, when coupled with the temporal discrimination arising from a kinetic asymmetry, leads to error correction and hence to the preservation of the evolutionarily dictated “order” during DNA replication. This model system thus provides a plausible scenario for the emergence of accurate information transfer across generations without enzymatic or energetic assistance from sources extraneous to the system, requiring only simple inorganic catalysts, such as metal ions, to catalyze covalent bond formation.

It is imperative that we answer our own question raised in the “Base Selection Energetics” subsection, within the “Introduction” section. What is the energy source for the error correction? We have claimed in the article that the irreversible covalent bond formation provides the necessary energy needed for error correction, although we did not quantitatively evaluate it. While the covalent linkage itself is effectively irreversible under physiological conditions—owing to the immediate hydrolysis of pyrophosphate, which suppresses the reverse reaction [98] – the energy released during this step constitutes the source of the non-equilibrium bias that favors accurate strand construction. In the main text, we treated this step as strictly irreversible, precluding a direct quantification of the corresponding free energy contribution. A more complete thermodynamic description-obtained by evaluating the total thermodynamic drive through the total entropy production rate at the nonequilibrium steady state [69]—is provided in the supplementary material (Section III).

The central aim of this article is to construct a highly simplified *model* that reproduces the essential features of error correcting systems, and to provide a deeper understanding of DNA replication fidelity. This necessitates a bare-bones approach, which led us to leave out many interactions that may have a bearing on the error correction ability, such as the nearest-neighbor thermodynamic interactions (stacking) between base pairs[99, 100], the dependence of error rates on local sequence characteristics[91], environmental effects on base pair bonding dynamics[101], and a multitude of other interactions that would be present in any highly evolved biological process. Even the assumption of the relationship between the two asymmetric cooperativity parameters, *α* and *β*, is made solely for simplicity, to reduce the number of parameters. Other assumptions, that only one location on the template is error-prone, that only two neighboring base pairs on either side affect the error rate, and that the covalent bonding is completely irreversible, are also made in a similar vein. Relaxing these assumptions may provide a more accurate picture closer to reality, but might diminish the results’ interpretability and the model’s pedagogical utility, in our view. Nevertheless, substantial improvements can be made to the model without jeopardizing its simplicity, for instance, by including sequence-dependent effects on error rates and matching them to experimentally observed sequence-dependent error rates[91]. The final arbiters of truth are experiments that can directly probe for the presence of asymmetric cooperativity in DNA replication, and examine its effects on error rates, possibly by modulating the cooperativity parameters through environmental perturbations or by any other means experimentally feasible. Identification of primordially plausible inorganic catalysts of covalent bond formation between nucleotides, such as montmorillonite clay[81, 102], would also strengthen our claims above. In conclusion, we have created a single-free-parameter theoretical model of error correction during DNA replication, demonstrating the possibility of utilizing the thermodynamic drive for self-replication itself for error correction, without needing any extraneous energy sources. We showed that the model reproduces several experimentally observed phenomena associated with error correction, and provides a possible explanation for the ability of thermodynamically passive polymerases to preferentially select correct base pairs. We noted that instantiations of the model are plausible in the primordial scenario because of its inherent simplicity. We showed that it is possible to create persistent order from free energy flow, provided we begin with appropriate symmetry-broken systems.

## Statements and Declarations

### Data availability

The code used to simulate the Markov chain dynamics is available in a private GitHub repository:https://github.com/KoushikGPhysics/New_Code_Non_Enzymatic_Error_Correction. Access will be provided upon request.

## Acknowledgments

Support for this work was provided by the Science & Engineering Research Board (SERB), Department of Science and Technology (DST), India, through a Core Research Grant with file no. CRG/2020/003555 and a MATRICS grant with file no. MTR/2022/000086.

## SUPPLEMENTARY MATERIAL

### I. Effect of the covalent bond formation between the 1^*st*^ and 2^*nd*^ nucleotides on the daughter strand construction fidelity

In the main text, our model incorporates “next-nucleotide effects”, whereby the formation of the (*m* + 1)^th^ base pair modulates covalent bond formation between the (*m* − 1)^th^ and *m*^th^ nucleotides in the daughter strand. Within the absorbing Markov chain framework, we focused specifically on covalent bond formation between the 2^nd^ and 3^rd^ nucleotides, assuming that covalent bonds between preceding nucleotides had already been established, while those between subsequent nucleotides have not yet formed. Although this simplified approach captures the essential kinetic discrimination, it remains important to verify that neglecting covalent bond formation between the 1^st^ and 2^nd^ nucleotides does not significantly affect the overall dynamics.

To examine the potential influence of covalent bond formation between the 1^*st*^ and 2^*nd*^ nucleotides on the dynamics, we extended the model to explicitly include this additional reaction. In this extended network, the covalent bond between the 1^*st*^ and 2^*nd*^ nucleotides forms according to the same rules applied for the bond between the 2^*nd*^ and 3^*rd*^ nucleotides, by requiring that the covalent bond between the (*m* − 1)^th^ and *m*^th^ nucleotides are established only after base pairing at the (*m* + 1)^th^ position. Fig.S1 presents a representative subset of transitions from the extended network, illustrating some of the reaction pathways.

We recomputed the transition rate matrix *Q* for this extended network and recalculated the accuracy ratio and stall quotient, as defined in Eqs.4 and 9, respectively. The results are shown in Fig.S2(a-c). In panel (a), we plot the accuracy ratio *η*_K+T_ as a function covalent bond formation rate *w*. Consistent with the Fig.7c, higher covalent bonding rates lead to improved error correction. Specifically, as *v* approaches and exceeds 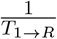, the covalent step between the 2^*nd*^ and 3^*rd*^ nucleotides effectively freezes in the third nucleotide, thereby amplifying any existing kinetic advantage and resulting in increase in the accuracy ratio.

In panel (b), we plot the accuracy ratio *η*_K+T_ as a function of effective base pairing drive Δ*G*_*C*_. As observed in Fig.8c, *η*_K+T_ displays a non-monotonic dependence on Δ*G*_*C*_, attaining the maximum value at intermediate value of Δ*G*_*C*_. Finally in panel (c), we plot the stall quotient *τ*_K+T_ as a function of effective base pairing drive Δ*G*_*C*_, which is also consistent with the Fig.9b.

Taken together, these findings indicate that the inclusion of covalent bond formation between the 1^*st*^ and 2^*nd*^ nucleotides in the daughter strand does not qualitatively modify the trends of kinetic or thermodynamic discrimination as reported in the main text. The extended network replicates the same characteristic dependencies of the accuracy ratio (*η*_K+T_) and stall quotient (*τ*_K+T_) on the covalent bond formation rate (*w*) and effective base-pairing drive (Δ*G*_*C*_). Consequently, limiting the analysis to covalent bond formation between the 2^nd^ and 3^rd^ nucleotides provides a physically faithful representation of the essential kinetic and thermodynamic discrimination that governs error correction in our model.

### II. Simplified Markov chain model of error correction with asymmetric cooperativity

Our aim here is to provide an approximate, analytically tractable Markov chain model for error correction, that captures the key observation that “covalent bond catalysis leads to error correction” in the main article. The variables and parameters used here are the same as defined in the Methods section of the main article. The analytical procedure followed here closely parallels the one used in [49].

We reduce the length of the template to *N* = 3, and assume that the construction of the daughter strand proceeds in a *strictly unidirectional manner*. This constraint reduces the dimensionality of the state space, facilitating analytical treatment. We characterize the dynamics in terms of a first-passage framework, wherein the principal quantity of interest is the first-passage time probability density function[60], which measures the temporal characteristics of the chain reaching a target state for the first time.

We use the backward (adjoint) master-equation approach to compute first-passage probability density. Let *f*_*i*→**R**_ denote the first-passage probability density for the process starting in state **i** at *t* = 0 to be first absorbed in **R** at time *t*. Similarly we define *f*_*i*→**W**_ for absorption into **W**. To solve them, we introduce **f**_**R**_(**t**) = (*F*_1→*R*_, *F*_2→*R*_, *F*_3→*R*_, *F*_4→*R*_, *F*_5→*R*_)^*T*^ and **f**_**W**_(**t**) = (*F*_1→*W*_, *F*_2→*W*_, *F*_3→*W*_, *F*_4→*W*_, *F*_5→*W*_)^*T*^. These satisfy the backward master equations:

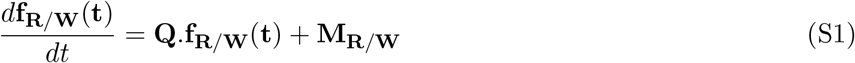

**Figure S1:**
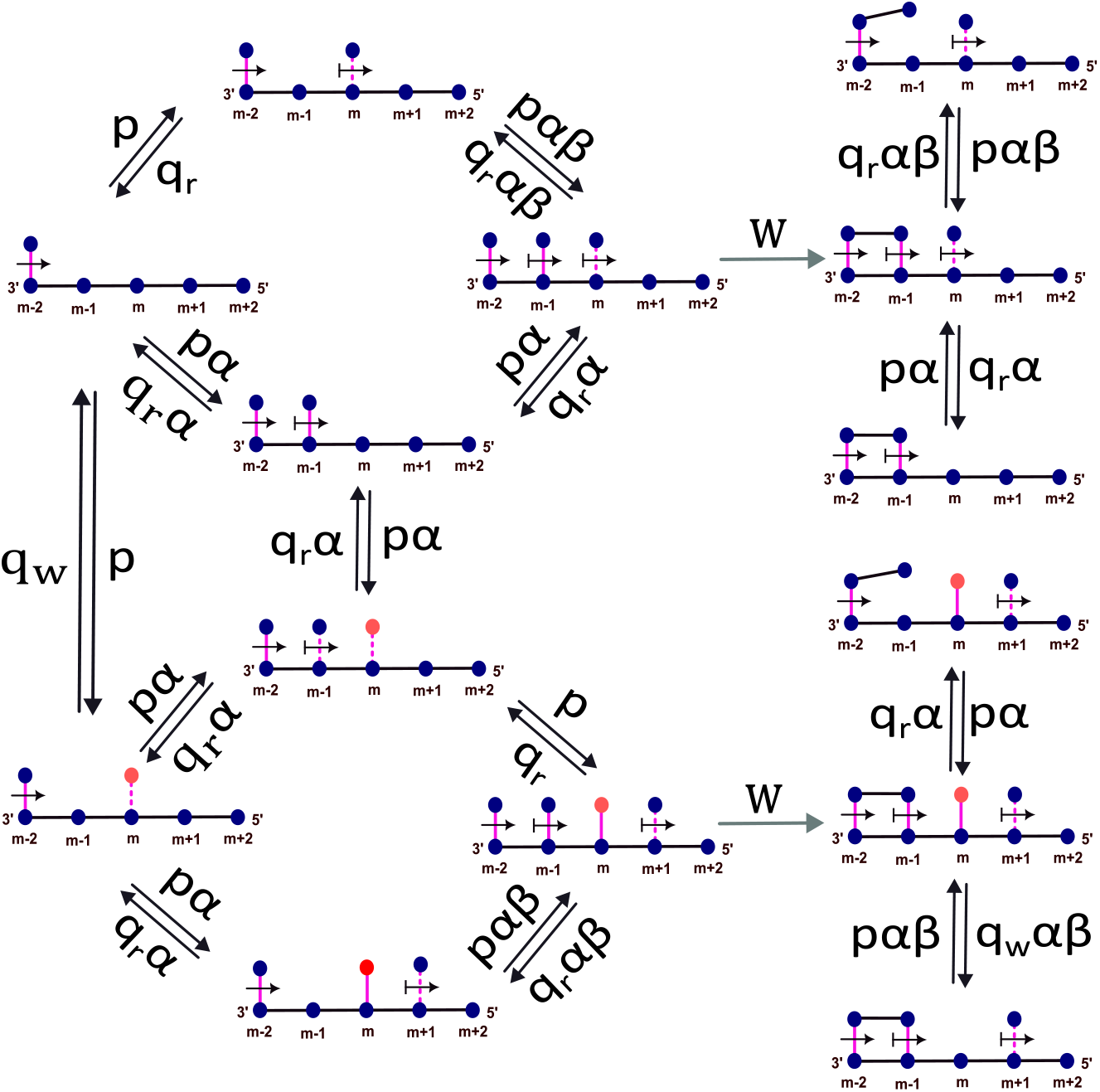
Simplified schematic of selected transitions from the covalent-bond formation network between the first and second base pairs (for *N* = 5). Covalent bond formation between nucleotides (*m* − 1) and *m* occurs only after base pairing at the (*m* + 1)^th^ position is established. For clarity, only a subset of representative transitions is shown; the full network includes additional states and paths.

**Figure S2:**
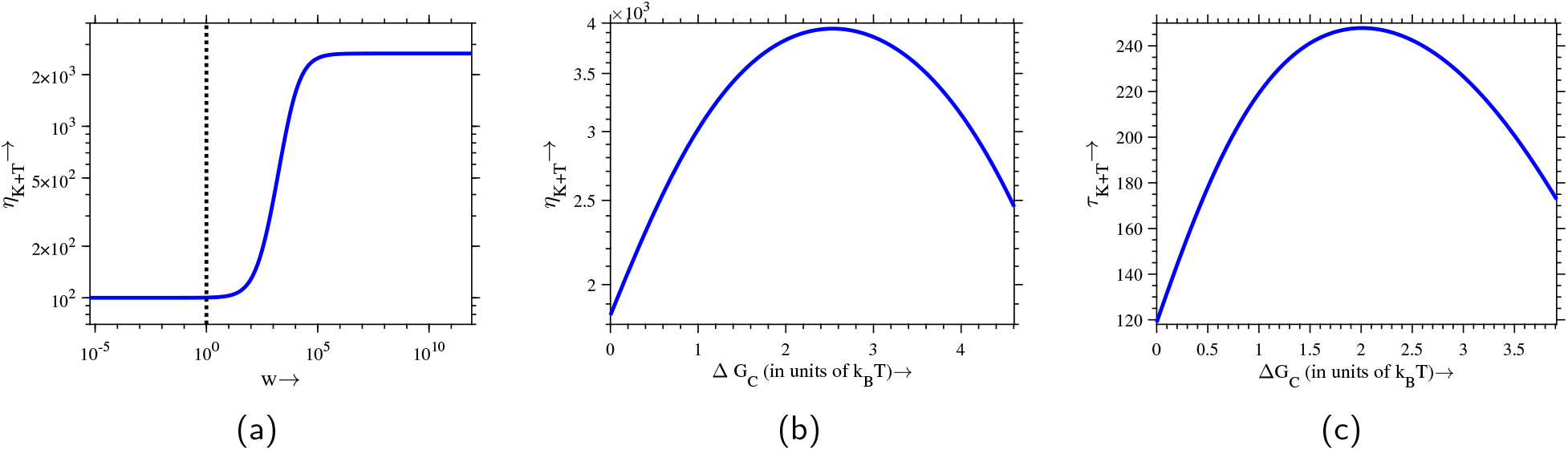
Accuracy ratio and stall quotient computed for the extended covalent-bond network. (a) Variation of the accuracy ratio *η*_K+T_ with the covalent-bond formation rate *w* (*p* = 2 sec^−1^, *q*_*r*_ = 1 sec^−1^, *q*_*w*_ = 100 sec^−1^, *α* = 5 *×* 10^3^). The plot The plot aligns with the kinetics–thermodynamics combined regime depicted in Fig.7, showing a sharp increase in accuracy once *v* approaches the base-pairing timescale (1*/T*_1→*R*_) i.e *w* ≈ 1. (b) Dependence of *η*_K+T_ on the effective base-pairing drive Δ*G*_*C*_ (*q*_*r*_ = 1 sec^−1^, *q*_*w*_ = 100 sec^−1^, *α* = 5 *×* 10^3^, *w* = 10^5^), showing the same non-monotonic trend observed in Fig.8c, where accuracy is maximized at an intermediate Δ*G*_*C*_. (c) Stall quotient *τ*_K+T_ as a function of Δ*G*_*C*_ (*q*_*r*_ = 1 sec^−1^, *q*_*w*_ = 100 sec^−1^, *α* = 5 *×* 10^3^, *w* = 10^5^)is also follows the same trend as in Fig.9b.

**Figure S3:**
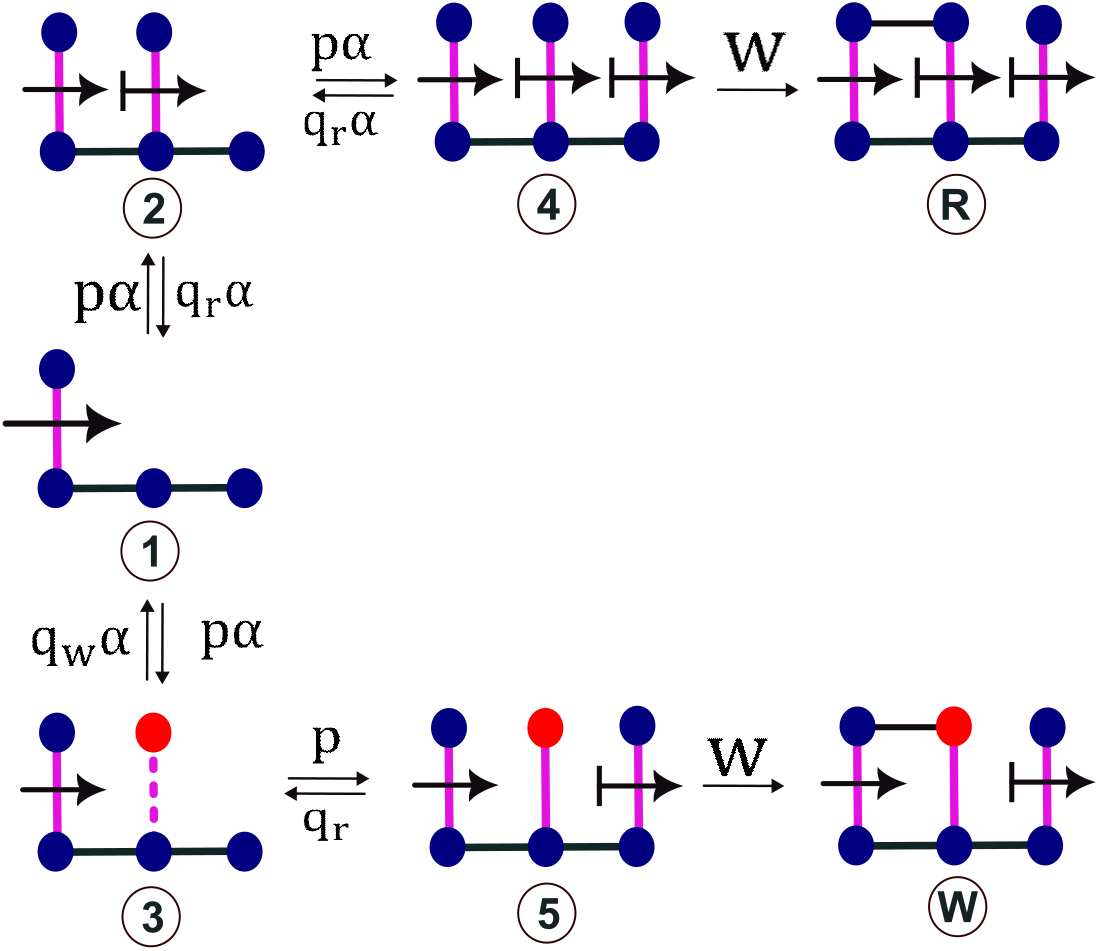
Minimal, approximate continuous-time Markov model for error correction with asymmetric cooperativity. Transient states represent intermediate hydrogen-bonded base-pairing configurations, while absorbing states correspond to the covalent incorporation of either a correct (R) or incorrect (W) nucleotide. The dynamics are governed by the base-pair formation rates(*p*), dissociation rates(*q*_*r*(*w*)_),the covalent locking rate *w*, and kinetic parameter *α* that modulates transition rates.

Here the term **Q.f**_**R***/***W**_(**t**) propagates probability through the hydrogen-bonded intermediates according to the usual adjoint chain dynamics, while the source vectors **M**_**R**_ and **M**_**W**_ inject probability corresponding to an immediate covalent-bond transition into **R** and **W**.

Now, from the network in Fig.S3 **M**_**R**_ = [0, 0, 0, *wδ*(*t*), 0]^*T*^ and **M**_**W**_ = [0, 0, 0, 0, *wδ*(*t*)]^*T*^, *δ*(*t*) being the Dirac delta function. The transition matrix is given by

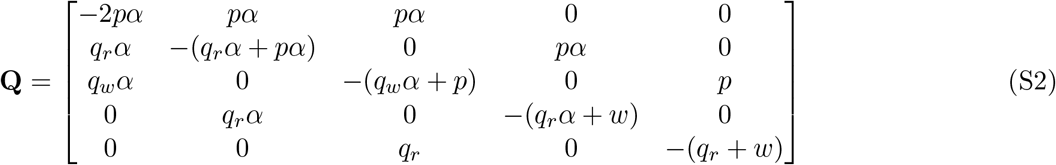

The Laplace transform fo *f*_*i*→**R***/***W**_(*t*) is defined as:

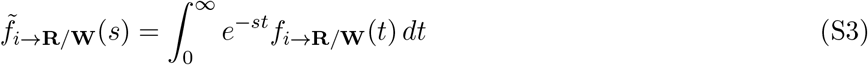

Taking the Laplace transform of the backward master Eq.S1, and **f**_**i**→**R***/***W**_(*t* = 0) = 0 for all (*I* ≠ **R***/***W**), we obtain:

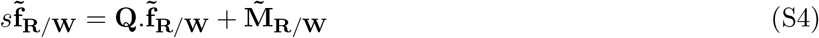

Here, 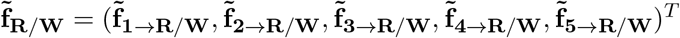, 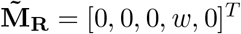 and 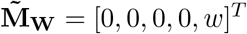. The Laplace-transformed first-passage probability densities 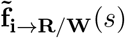 are then obtained by solving the algebraic set of Eqs.S4. The *splitting probabilities*, which describe the likelihood of eventually reaching either absorbing state, are obtained by evaluating 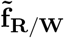 at *s* = 0:

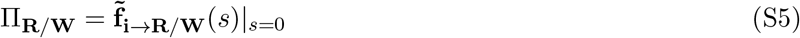

From the analytical solution of the model, the ratio of splitting probabilities is given by:

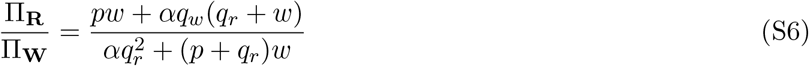

This expression exhibits two distinct limiting behaviors:

**Thermodynamic regime**, w is low:

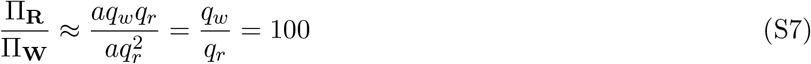

Therefore, the accuracy of the daughter strand construction is purely controlled by the base-pair dissociation rates. The system has sufficient time to equilibrate before covalent locking occurs, so accuracy is entirely thermodynamically controlled.

**Kinetic regime**, w is high and *p << αq*_*w*_:

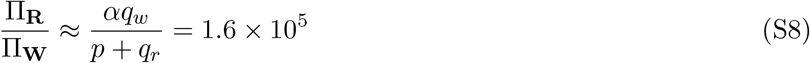

Here, rapid covalent bond formation prevents equilibration and allows kinetic discrimination to persist indefinitely.

### III. Thermodynamic analysis of the accuracy-dissipation trade-off

Extending the analysis presented in Section 4.III, we quantify the overall thermodynamic drive associated with daughter-strand construction by evaluating the total entropy production rate (EPR) of the reaction network, which now includes covalent bond formation. In this formulation, the EPR incorporates all energetic contributions accumulated along both correct and incorrect incorporation pathways, thereby providing a measure of the thermodynamic cost associated with strand extension. The entropy production rate (EPR) of the entire reaction network can be computed using the Schnakenberg formulation [103] for Markov processes:

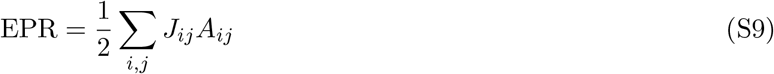

where *J*_*ij*_ = *P*_*i*_*Q*_*ij*_ − *P*_*j*_*Q*_*ji*_ denotes the net steady-state flux between states *i* and *j*, and 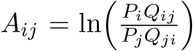 is the corresponding thermodynamic affinity. The probabilities *P*_*i*_ are the steady-state occupancies of network states, and *Q*_*ij*_ are the transition rates defined in the master equation 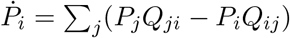.

The results of this analysis are shown in Fig.S4, where we observe the same qualitative dependence of *η* on the EPR as observed in Fig.8, where *η* was plotted as a function of Δ*G*_C_. This observation points to the existence of an optimal dissipation regime, beyond which additional thermodynamic drive no longer improves the accuracy.

**Figure S4:**
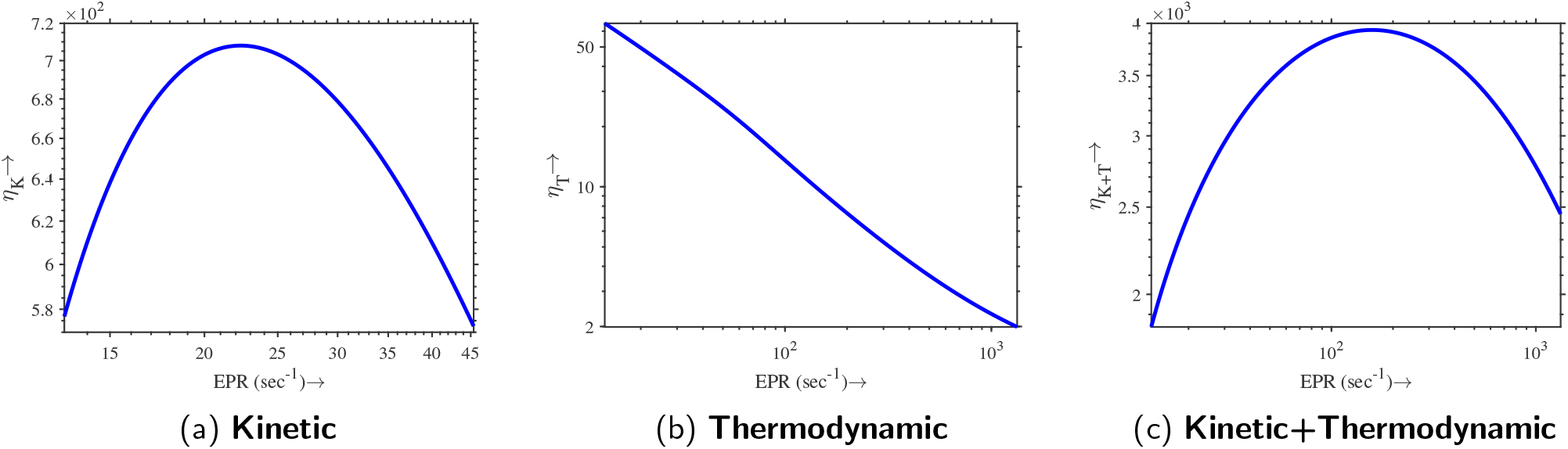
Illustration of the accuracy ratio as a function of the entropy production rate (EPR) across three distinct regimes. Panels (a) and (b) correspond to the purely kinetic and purely thermodynamic discrimination, respectively, while panel (c) depicts the combined (kinetic + thermodynamic) discrimination scenario. All three plots exhibit the same qualitative trends observed in Fig.8.

### IV. Determination of *α* from experimentally determined ratio of correct and incorrect base-pair dissociation rates

To set an appropriate value for the kinetic parameter *α*, we make use of the experimentally determined ratio of rates of dissociation of incorrect to correct base pairs, of the order of 10^3^, in [72], for T7 DNA polymerase. This ratio is simply MFPT_11100→11000_*/*MFPT_11200→11000_, within our model. In Fig.S5, we show the variation of this ratio as a function of *α*, from which we derive a value of 5 *×* 10^3^ for *α*.

**Figure S5:**
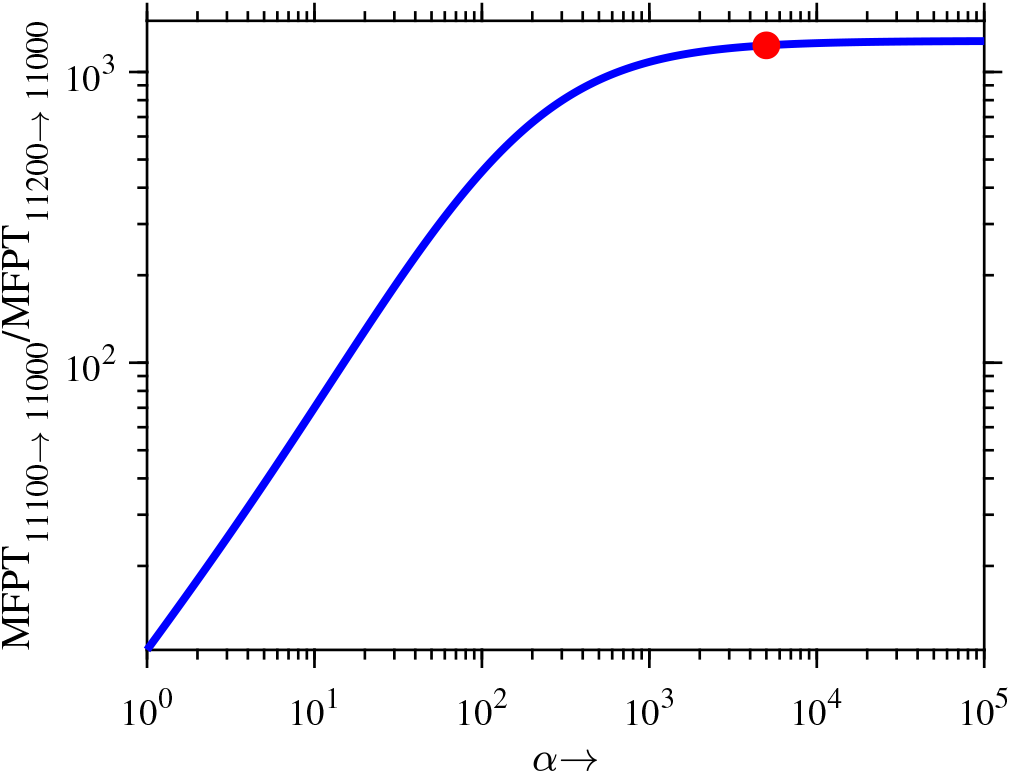
Variation of the ratio of mean first passage times (MFPT),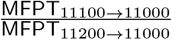, as a function of the asymmetric cooperativity parameter *α*. Here, MFPT _11100*/*11200→11000_ denotes the MFPT for the system starting from either the 11100 or 11200 state to reach the state 11000. As *α* increases, MFPT_11100→11000_ becomes progressively larger than MFPT_11200→11000_, and their ratio approaches ∼ 10^3^ at *α* ∼ 5 *×* 10^3^.

### V. Comparison of absorbing and non-absorbing Markov chain models

To provide a basis for considering the absorbing Markov chain framework for our model, we compared the non-absorbing and absorbing Markov formulations of daughter strand construction. The details of this comparison are presented below.

**Figure S6:**
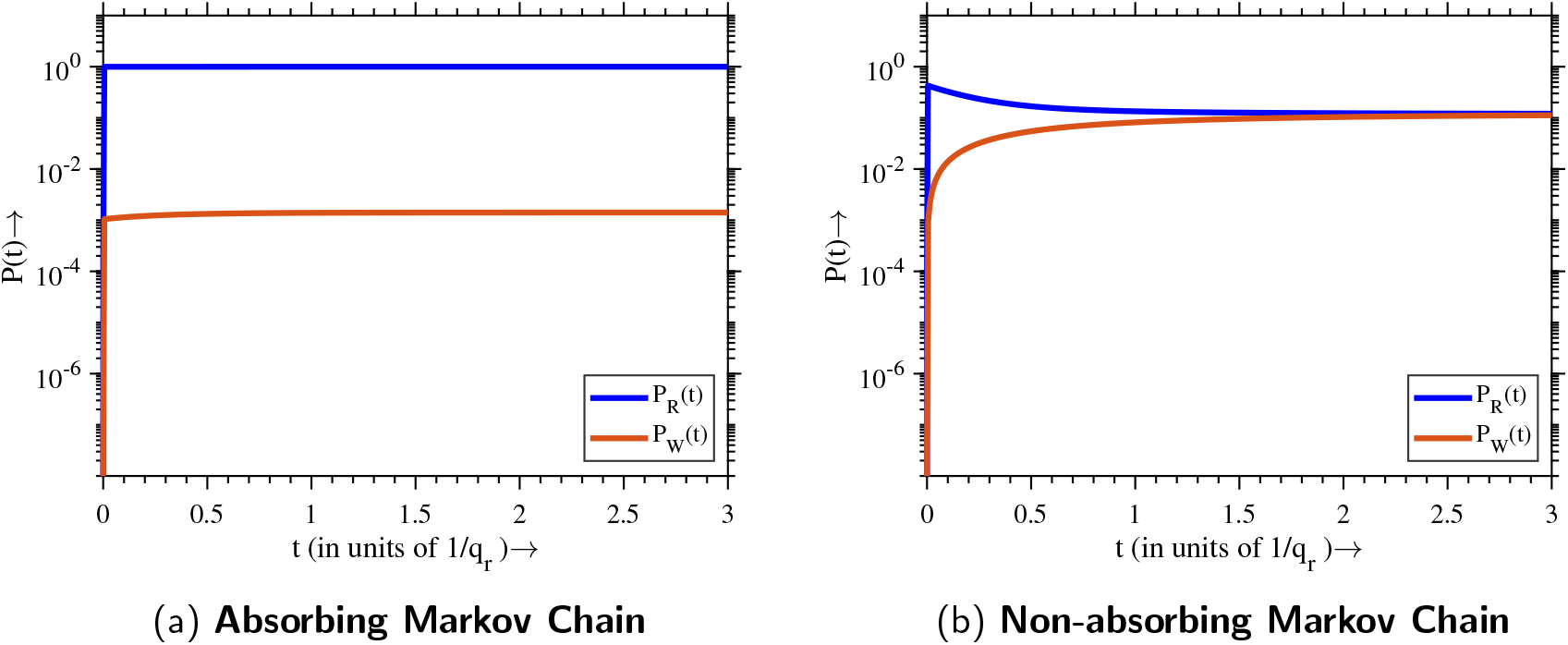
Time evolution of the occupation probabilities for the correct state 11111 and the incorrect state 11211, denoted by *P*_*R*_(*t*) and *P*_*W*_ (*t*), respectively. (a) In the absorbing Markov chain framework, kinetic discrimination persists at steady state due to the irreversible nature of the final state, which drives the system out of equilibrium. The probabilities of all intermediate states go to zero at steady state. (b) In the non-absorbing Markov chain, the system evolves toward equilibrium, and kinetic discrimination vanishes at steady state as detailed balance is satisfied. The probabilities of intermediate states (not shown) are non-zero.

#### i. Kinetic discrimination leads to error correction only under irreversible conditions

Here, we study the role of kinetic discrimination alone by examining how it operates under two different modeling approaches. Specifically, we compare an absorbing and a non-absorbing Markov chain model to assess whether kinetic discrimination alone, without any thermodynamic bias, i.e., with *q*_*r*_ = *q*_*w*_ = 1 sec^−1^, can sustain daughter-strand fidelity in each case.

In Fig.S6, we plot the time-evolution of the occupation probability of the two final states 11111 and 11211, denoted by *P*_*R*_(*t*) and *P*_*w*_(*t*), respectively. These trajectories shows the fundamental differences between the absorbing and non-absorbing models in the long-time limit. In the absorbing Markov chain (Fig.S6a), kinetic discrimination persists even at steady state. This is because of the presence of an absorbing final state, which makes the system inherently out of equilibrium, thereby violating the detailed balance condition. As a result, kinetic asymmetries continue to influence the dynamics, due to the persistent flow of probabilities from the initial to the final states, allowing for sustained generation of accurate daughter strands. In contrast, in the non-absorbing Markov chain (Fig.S6b), kinetic discrimination vanishes at steady state. Since the discrimination is based on the time difference between the correct and incorrect daughter strands, and since the steady state is time-independent, kinetic discrimination becomes ineffective. The local detailed balance [104] holds and entropy production becomes zero, and the system reaches equilibrium, despite the presence of a thermodynamic drive, suggesting that the system is unable to absorb the energy from the drive due to the possibility of reverse transitions.

This observation indicates that kinetic discrimination can only be sustained either during the transient phase of a non-absorbing Markov chain or by rendering the final-state transition effectively irreversible, thereby transforming the system into an absorbing Markov chain. In this study, we adopt the latter approach by employing an absorbing Markov chain model.

#### ii. Kinetic+thermodynamic discrimination improves error correction in absorbing Markov chain

Now we proceed to examine how the integration of kinetic and thermodynamic discrimination improves error correction when the absorbing and non-absorbing Markov chain modeling approaches are used. To incorporate thermodynamic discrimination alongside kinetic discrimination, we consider the dissociation rate of the correct base pair as *q*_*r*_ = 1 sec^−1^ and that of the incorrect base pair as *q*_*w*_ = 100 sec^−1^, hundred times larger than *q*_*r*_ (Methods).

**Figure S7:**
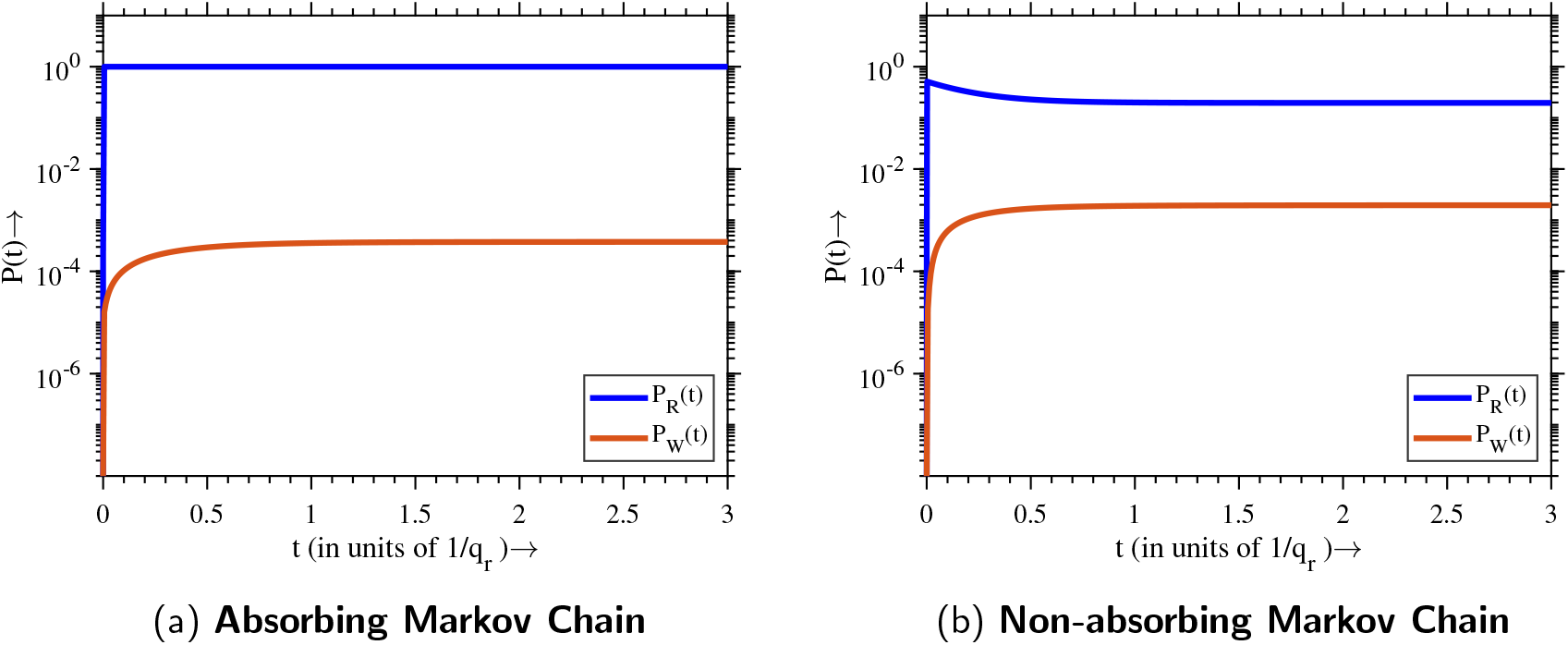
Time evolution of the occupation probabilities for the correct state 11111 and the erroneous state 11211, denoted by *P*_*R*_(*t*) and *P*_*W*_ (*t*), respectively, under combined kinetic and thermodynamic discrimination. (a) In the absorbing Markov chain, the system remains out of equilibrium, allowing both kinetic and thermodynamic discrimination to contribute to improved accuracy. (b) In the non-absorbing Markov chain, the system reaches equilibrium at long times, and only thermodynamic discrimination governs the steady-state accuracy, which is lower than the accuracy obtained in the absorbing case.

As shown in Fig.S7a, in the absorbing Markov chain approach, combining kinetic and thermodynamic discrimination improves the accuracy ratio *η*_K+T_ beyond what either mechanism can achieve on its own. The absorbing final state keeps the system out of equilibrium, allowing kinetic differences to continue influencing the dynamics and work together with thermodynamic bias to enhance fidelity. In contrast, in the non-absorbing Markov chain, Fig.S7b, the system eventually reaches equilibrium, where kinetic discrimination no longer plays a role due to detailed balance. As a result, in this case, the accuracy ratio *η*_K+T_ is limited by thermodynamic discrimination alone.

### VI. Dependence of accuracy ratio and stall quotient on template length N

**Figure S8:**
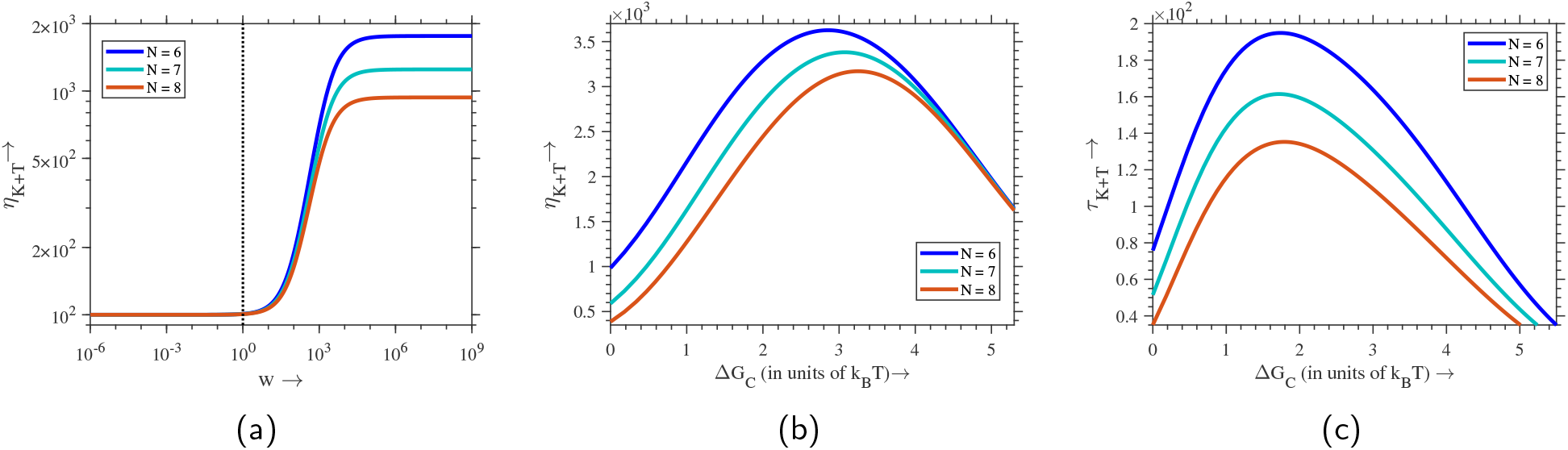
Variation of accuracy ratio (*η*_K+T_) and the stall quotient (*τ*_K+T_) with the template sequence length *N*. This demonstrates that extending the analysis to longer templates produces the same qualitative behavior. The curves for *η*_K+T_ and *τ*_K+T_ maintain the same characteristic shape across the three *N* values tested here.

## References

[1] Charles Darwin. On the origin of species: A facsimile of the first edition. Harvard University Press, 1964.

[2] William Bateson and Gregor Mendel. Mendel’s principles of heredity. Courier Corporation, 2013.

[3] John J Hopfield. Kinetic proofreading: a new mechanism for reducing errors in biosynthetic processes requiring high specificity. Proceedings of the National Academy of Sciences, 71(10):4135–4139, 1974.

[4] Jacques Ninio. Kinetic amplification of enzyme discrimination. Biochimie, 57(5):587–595, 1975.

[5] Alexandra M De Paz, Thaddeus R Cybulski, Adam H Marblestone, Bradley M Zamft, George M Church, Edward S Boyden, Konrad P Kording, and Keith E J Tyo. High-resolution mapping of DNA polymerase fidelity using nucleotide imbalances and next-generation sequencing. Nucleic acids research, 46(13):e78– e78, 2018.

[6] Kingsley L Wong and Juewen Liu. Factors and methods to modulate DNA hybridization kinetics. Biotechnology journal, 16(11):2000338, 2021.

[7] Rais A Ganai and Erik Johansson. DNA replication—a matter of fidelity. Molecular cell, 62(5):745–755, 2016.

[8] Manfred Eigen. Selforganization of matter and the evolution of biological macromolecules. Naturwissenschaften, 58:465–523, 1971.

[9] Manfred Eigen and Peter Schuster. The hypercycle: a principle of natural self-organization. Springer Science & Business Media, 2012.

[10] Josep Sardanyés, Celia Perales, Esteban Domingo, and Santiago F Elena. Quasispecies theory and emerging viruses: challenges and applications. npj Viruses, 2(1):54, 2024.

[11] JJ Hopfield, T Yamane, V Yue, and SM Coutts. Direct experimental evidence for kinetic proofreading in amino acylation of tRNAIle. Proceedings of the National Academy of Sciences, 73(4):1164–1168, 1976.

[12] Arvind Murugan, David A Huse, and Stanislas Leibler. Speed, dissipation, and error in kinetic proof-reading. Proceedings of the National Academy of Sciences, 109(30):12034–12039, 2012.

[13] T Yamane and JJ Hopfield. Experimental evidence for kinetic proofreading in the aminoacylation of tRNA by synthetase. Proceedings of the National Academy of Sciences, 74(6):2246–2250, 1977.

[14] Scott C Blanchard, Ruben L Gonzalez Jr, Harold D Kim, Steven Chu, and Joseph D Puglisi. tRNA selection and kinetic proofreading in translation. Nature structural & molecular biology, 11(10):1008– 1014, 2004.

[15] Joel D Mallory, Anatoly B Kolomeisky, and Oleg A Igoshin. Trade-offs between error, speed, noise, and energy dissipation in biological processes with proofreading. The Journal of Physical Chemistry B, 123(22):4718–4725, 2019.

[16] Qiwei Yu, Joel D Mallory, Anatoly B Kolomeisky, Jiqiang Ling, and Oleg A Igoshin. Trade-offs between speed, accuracy, and dissipation in tRNAIle aminoacylation. The journal of physical chemistry letters, 11(10):4001–4007, 2020.

[17] Robert A Beckman and Lawrence A Loeb. Multi-stage proofreading in DNA replication. Quarterly reviews of biophysics, 26(3):225–331, 1993.

[18] Scott D McCulloch and Thomas A Kunkel. The fidelity of DNA synthesis by eukaryotic replicative and translesion synthesis polymerases. Cell research, 18(1):148–161, 2008.

[19] Harrison Echols and Myron F Goodman. Fidelity mechanisms in DNA replication. Annual review of biochemistry, 60(1):477–511, 1991.

[20] Myron F Goodman, Steven Creighton, Linda B Bloom, John Petruska, and Thomas A Kunkel. Biochemical basis of DNA replication fidelity. Critical reviews in biochemistry and molecular biology, 28(2):83–126, 1993.

[21] Andrew C Olson, Jennifer N Patro, Milan Urban, and Robert D Kuchta. The energetic difference between synthesis of correct and incorrect base pairs accounts for highly accurate DNA replication. Journal of the american chemical society, 135(4):1205–1208, 2013.

[22] Shina CL Kamerlin, Pankaz K Sharma, Ram B Prasad, and Arieh Warshel. Why nature really chose phosphate. Quarterly reviews of biophysics, 46(1):1–132, 2013.

[23] Keriann Oertell, Emily M Harcourt, Michael G Mohsen, John Petruska, Eric T Kool, and Myron F Goodman. Kinetic selection vs. free energy of DNA base pairing in control of polymerase fidelity. Proceedings of the National Academy of Sciences, 113(16):E2277–E2285, 2016.

[24] Thomas A Kunkel. Evolving views of dna replication (in) fidelity. In Cold Spring Harbor symposia on quantitative biology, volume 74, pages 91–101. Cold Spring Harbor Laboratory Press, 2009.

[25] John Petruska, Lawrence C Sowers, and Myron F Goodman. Comparison of nucleotide interactions in water, proteins, and vacuum: model for DNA polymerase fidelity. Proceedings of the National Academy of Sciences, 83(6):1559–1562, 1986.

[26] Michael S Boosalis, J Petruska, and MF Goodman. DNA polymerase insertion fidelity. Gel assay for site-specific kinetics. Journal of Biological Chemistry, 262(30):14689–14696, 1987.

[27] David Andrieux and Pierre Gaspard. Nonequilibrium generation of information in copolymerization processes. Proceedings of the National Academy of Sciences, 105(28):9516–9521, 2008.

[28] Paul G Higgs and Niles Lehman. The RNA World: molecular cooperation at the origins of life. Nature Reviews Genetics, 16(1):7–17, 2015.

[29] Gerald F Joyce and Jack W Szostak. Protocells and rna self-replication. Cold Spring Harbor Perspectives in Biology, 10(9):a034801, 2018.

[30] Patrizia Hagenbuch, Eric Kervio, Annette Hochgesand, Ulrich Plutowski, and Clemens Richert. Chemical primer extension: efficiently determining single nucleotides in DNA. Angewandte Chemie, 117(40):6746– 6750, 2005.

[31] Jianyang Han, Eric Kervio, and Clemens Richert. High Fidelity Enzyme-Free Primer Extension with an Ethynylpyridone Thymidine Analog. Chemistry–A European Journal, 27(64):15918–15921, 2021.

[32] Tobias Göppel, Benedikt Obermayer, Irene A Chen, and Ulrich Gerland. A kinetic error filtering mechanism for enzyme-free copying of nucleic acid sequences. bioRxiv, pages 2021–08, 2021.

[33] Yoshiya J Matsubara, Nobuto Takeuchi, and Kunihiko Kaneko. Avoidance of error catastrophe via proofreading innate to template-directed polymerization. Physical Review Research, 5(1):013170, 2023.

[34] Rakesh Mukherjee, Aditya Sengar, Javier Cabello-García, and Thomas E Ouldridge. Kinetic proofreading can enhance specificity in a nonenzymatic DNA strand displacement network. Journal of the American Chemical Society, 146(28):18916–18926, 2024.

[35] Vahe Galstyan, Kabir Husain, Fangzhou Xiao, Arvind Murugan, and Rob Phillips. Proofreading through spatial gradients. Elife, 9:e60415, 2020.

[36] Anthonie WJ Muller and Dirk Schulze-Makuch. Thermal energy and the origin of life. Origins of Life and Evolution of Biospheres, 36:177–189, 2006.

[37] Laurent Boiteau and Robert Pascal. Energy sources, self-organization, and the origin of life. Origins of Life and Evolution of Biospheres, 41(1):23–33, 2011.

[38] Kepa Ruiz-Mirazo, Carlos Briones, and Andres de la Escosura. Prebiotic systems chemistry: new perspectives for the origins of life. Chemical reviews, 114(1):285–366, 2014.

[39] Ilya Prigogine and Grégoire Nicolis. Self-organisation in nonequilibrium systems: towards a dynamics of complexity. Bifurcation analysis, pages 3–12, 1985.

[40] Todd R Gingrich, Jordan M Horowitz, Nikolay Perunov, and Jeremy L England. Dissipation bounds all steady-state current fluctuations. Physical review letters, 116(12):120601, 2016.

[41] Jordan M Horowitz, Kevin Zhou, and Jeremy L England. Minimum energetic cost to maintain a target nonequilibrium state. Physical Review E, 95(4):042102, 2017.

[42] Jordan M Horowitz and Jeremy L England. Spontaneous fine-tuning to environment in many-species chemical reaction networks. Proceedings of the National Academy of Sciences, 114(29):7565–7570, 2017.

[43] Hemachander Subramanian and Robert A Gatenby. Evolutionary advantage of directional symmetry breaking in self-replicating polymers. Journal of theoretical biology, 446:128–136, 2018.

[44] Hemachander Subramanian and Robert A Gatenby. Evolutionary advantage of anti-parallel strand orientation of duplex DNA. Scientific Reports, 10(1):9883, 2020.

[45] Sudha Rajamani, Justin K Ichida, Tibor Antal, Douglas A Treco, Kevin Leu, Martin A Nowak, Jack W Szostak, and Irene A Chen. Effect of stalling after mismatches on the error catastrophe in nonenzymatic nucleic acid replication. Journal of the American Chemical Society, 132(16):5880–5885, 2010.

[46] Kevin Leu, Eric Kervio, Benedikt Obermayer, Rebecca M Turk-MacLeod, Caterina Yuan, Jesus-Mario Luevano Jr, Eric Chen, Ulrich Gerland, Clemens Richert, and Irene A Chen. Cascade of reduced speed and accuracy after errors in enzyme-free copying of nucleic acid sequences. Journal of the American Chemical Society, 135(1):354–366, 2013.

[47] Tobias Göppel, Joachim H Rosenberger, Bernhard Altaner, and Ulrich Gerland. Thermodynamic and kinetic sequence selection in enzyme-free polymer self-assembly inside a non-equilibrium RNA reactor. Life, 12(4):567, 2022.

[48] Riccardo Ravasio, Kabir Husain, Constantine G Evans, Rob Phillips, Marco Ribezzi, Jack W Szostak, and Arvind Murugan. A minimal scenario for the origin of non-equilibrium order. arXiv preprint 2405.10911, 2024.

[49] Kinshuk Banerjee, Anatoly B Kolomeisky, and Oleg A Igoshin. Elucidating interplay of speed and accuracy in biological error correction. Proceedings of the National Academy of Sciences, 114(20):5183– 5188, 2017.

[50] Jenny M Poulton, Pieter Rein Ten Wolde, and Thomas E Ouldridge. Nonequilibrium correlations in minimal dynamical models of polymer copying. Proceedings of the National Academy of Sciences, 116(6):1946– 1951, 2019.

[51] Vahe Galstyan and Rob Phillips. Allostery and kinetic proofreading. The Journal of Physical Chemistry B, 123(51):10990–11002, 2019.

[52] Rafael Fernandez-Leiro, Julian Conrad, Ji-Chun Yang, Stefan MV Freund, Sjors HW Scheres, and Meindert H Lamers. Self-correcting mismatches during high-fidelity DNA replication. Nature structural & molecular biology, 24(2):140–143, 2017.

[53] Thomas A Kunkel, RM Schaaper, RA Beckman, and LA Loeb. On the fidelity of DNA replication. Effect of the next nucleotide on proofreading. Journal of Biological Chemistry, 256(19):9883–9889, 1981.

[54] Toon Swings, Bram Van den Bergh, Sander Wuyts, Eline Oeyen, Karin Voordeckers, Kevin J Verstrepen, Maarten Fauvart, Natalie Verstraeten, and Jan Michiels. Adaptive tuning of mutation rates allows fast response to lethal stress in Escherichia coli. Elife, 6:e22939, 2017.

[55] Troy G Hammerstrom, Kathryn Beabout, Thomas P Clements, Gerda Saxer, and Yousif Shamoo. Acinetobacter baumannii repeatedly evolves a hypermutator phenotype in response to tigecycline that effectively surveys evolutionary trajectories to resistance. PloS one, 10(10):e0140489, 2015.

[56] Jithesh Kottur and Deepak T Nair. Pyrophosphate hydrolysis is an intrinsic and critical step of the DNA synthesis reaction. Nucleic Acids Research, 46(12):5875–5885, 2018.

[57] Raymond Dean Astumian. Kinetic Asymmetry and directionality of nonequilibrium molecular systems. Angewandte Chemie, 136(9):e202306569, 2024.

[58] Lei Tang, Luis A Navarro Jr, Ashutosh Chilkoti, and Stefan Zauscher. Angewandte Chemie, 129(24):6882–6886, 2017.

[59] Chengjie Zhang, Hizar Subthain, Fei Guo, Peng Fang, Shanmin Zheng, Mengzhe Shen, Xianger Jiang, Zhengquan Gao, Chunxiao Meng, Shengying Li, et al. Terminal deoxynucleotidyl transferase: Properties and applications. Engineering Microbiology, page 100179, 2024.

[60] NG Van Kampen. Stochastic Processes in Physics and Chemistry. North-Holland Publishing Co, 1992.

[61] William J Anderson. Continuous-time Markov chains: An applications-oriented approach. Springer Science & Business Media, 2012.

[62] Oliver Ibe. Markov processes for stochastic modeling. Newnes, 2013.

[63] Robert P Dobrow. Introduction to stochastic processes with R. John Wiley & Sons, 2016.

[64] Peter M Burgers. Solution to the 50-year-old Okazaki-fragment problem. Proceedings of the National Academy of Sciences, 116(9):3358–3360, 2019.

[65] Sean P Fagan, Purba Mukherjee, William J Jaremko, Rachel Nelson-Rigg, Ryan C Wilson, Tyler L Dangerfield, Kenneth A Johnson, Indrajit Lahiri, and Janice D Pata. Pyrophosphate release acts as a kinetic checkpoint during high-fidelity DNA replication by the Staphylococcus aureus replicative polymerase PolC. Nucleic Acids Research, 49(14):8324–8338, 2021.

[66] Travis Walton, Wen Zhang, Li Li, Chun Pong Tam, and Jack W Szostak. The mechanism of nonenzymatic template copying with imidazole-activated nucleotides. Angewandte Chemie International Edition, 58(32):10812–10819, 2019.

[67] Stephanie R Vogel, Christopher Deck, and Clemens Richert. Accelerating chemical replication steps of RNA involving activated ribonucleotides and downstream-binding elements. Chemical Communications, (39):4922–4924, 2005.

[68] Paul C Bressloff. Search processes with stochastic resetting and multiple targets. Physical Review E, 102(2):022115, 2020.

[69] Kinshuk Banerjee, Biswajit Das, and Gautam Gangopadhyay. The guiding role of dissipation in kinetic proofreading networks: Implications for protein synthesis. The Journal of Chemical Physics, 152(11), 2020.

[70] Stefan Howorka, Liviu Movileanu, Orit Braha, and Hagan Bayley. Kinetics of duplex formation for individual DNA strands within a single protein nanopore. Proceedings of the National Academy of Sciences, 98(23):12996–13001, 2001.

[71] Elizaveta Guseva, Ronald N Zuckermann, and Ken A Dill. Foldamer hypothesis for the growth and sequence differentiation of prebiotic polymers. Proceedings of the National Academy of Sciences, 114(36):E7460–E7468, 2017.

[72] Yu-Chih Tsai and Kenneth A Johnson. A new paradigm for DNA polymerase specificity. Biochemistry, 45(32):9675–9687, 2006.

[73] Pablo Sartori and Simone Pigolotti. Kinetic versus energetic discrimination in biological copying. Physical review letters, 110(18):188101, 2013.

[74] Thomas E Ouldridge, Petr Šulc, Flavio Romano, Jonathan PK Doye, and Ard A Louis. DNA hybridization kinetics: zippering, internal displacement and sequence dependence. Nucleic acids research, 41(19):8886– 8895, 2013.

[75] Sophie Hertel, Richard E Spinney, Stephanie Y Xu, Thomas E Ouldridge, Richard G Morris, and Lawrence K Lee. The stability and number of nucleating interactions determine DNA hybridization rates in the absence of secondary structure. Nucleic Acids Research, 50(14):7829–7841, 2022.

[76] Marco Todisco and Jack W Szostak. Hybridization kinetics of out-of-equilibrium mixtures of short RNA oligonucleotides. Nucleic Acids Research, 50(17):9647–9662, 2022.

[77] Isaac Wong, Smita S Patel, and Kenneth A Johnson. An induced-fit kinetic mechanism for DNA replication fidelity: direct measurement by single-turnover kinetics. Biochemistry, 30(2):526–537, 1991.

[78] John W Brandis, Sydney G Edwards, and Kenneth A Johnson. Slow rate of phosphodiester bond formation accounts for the strong bias that Taq DNA polymerase shows against 2 ‘, 3 ‘-dideoxynucleotide terminators. Biochemistry, 35(7):2189–2200, 1996.

[79] Bret D Freudenthal, William A Beard, David D Shock, and Samuel H Wilson. Observing a DNA polymerase choose right from wrong. Cell, 154(1):157–168, 2013.

[80] Anuraag Aithal, Shikha Dagar, and Sudha Rajamani. Metals in prebiotic catalysis: a possible evolutionary pathway for the emergence of metalloproteins. ACS omega, 8(6):5197–5208, 2023.

[81] James P Ferris, Aubrey R Hill Jr, Rihe Liu, and Leslie E Orgel. Synthesis of long prebiotic oligomers on mineral surfaces. Nature, 381(6577):59–61, 1996.

[82] Arvind Murugan, David A Huse, and Stanislas Leibler. Discriminatory proofreading regimes in nonequilibrium systems. Physical Review X, 4(2):021016, 2014.

[83] Chunhong Long and Jin Yu. Balancing Non-Equilibrium driving with nucleotide selectivity at kinetic checkpoints in polymerase fidelity control. Entropy, 20(4):306, 2018.

[84] Brian Munsky, Ilya Nemenman, and Golan Bel. Specificity and completion time distributions of biochemical processes. The Journal of chemical physics, 131(23), 2009.

[85] Riccardo Rao and Luca Peliti. Thermodynamics of accuracy in kinetic proofreading: dissipation and efficiency trade-offs. Journal of Statistical Mechanics: Theory and Experiment, 2015(6):P06001, 2015.

[86] Kristian Le Vay, Elia Salibi, Emilie Y Song, and Hannes Mutschler. Nucleic acid catalysis under potential prebiotic conditions. Chemistry–An Asian Journal, 15(2):214–230, 2020.

[87] Oliver J Rando and Kevin J Verstrepen. Timescales of genetic and epigenetic inheritance. Cell, 128(4):655–668, 2007.

[88] Jeffrey H Chuang and Hao Li. Functional bias and spatial organization of genes in mutational hot and cold regions in the human genome. PLoS biology, 2(2):e29, 2004.

[89] Marjan W Van Der Woude and Andreas J B”aumler. Phase and antigenic variation in bacteria. Clinical microbiology reviews, 17(3):581–611, 2004.

[90] J David Barry and Richard McCulloch. Antigenic variation in trypanosomes: enhanced phenotypic variation in a eukaryotic parasite. Advances in Parasitology, 49:1–70, 2001.

[91] Eric Kervio, Annette Hochgesand, Ulrich E Steiner, and Clemens Richert. Templating efficiency of naked DNA. Proceedings of the National Academy of Sciences, 107(27):12074–12079, 2010.

[92] Eörs Szathmáry. The origin of replicators and reproducers. Philosophical Transactions of the Royal Society B: Biological Sciences, 361(1474):1761–1776, 2006.

[93] Thomas A Kunkel. DNA replication fidelity. Journal of Biological Chemistry, 279(17):16895–16898, 2004.

[94] William A Beard and Samuel H Wilson. Structural insights into the origins of DNA polymerase fidelity. Structure, 11(5):489–496, 2003.

[95] Thomas W Traut. Physiological concentrations of purines and pyrimidines. Molecular and cellular biochemistry, 140:1–22, 1994.

[96] Erwin Schrödinger. What is life?: With mind and matter and autobiographical sketches. Cambridge university press, 1992.

[97] Philip W Anderson and Daniel L Stein. Broken symmetry, emergent properties, dissipative structures, life: Are they related? In Basic notions of condensed matter physics, pages 263–277. CRC Press, 2018.

[98] Jessica LE Wimmer, Karl Kleinermanns, and William F Martin. Pyrophosphate and irreversibility in evolution, or why PPi is not an energy currency and why nature chose triphosphates. Frontiers in microbiology, 12:759359, 2021.

[99] John Petruska and MF Goodman. Influence of neighboring bases on DNA polymerase insertion and proofreading fidelity. Journal of Biological Chemistry, 260(12):7533–7539, 1985.

[100] Peter Yakovchuk, Ekaterina Protozanova, and Maxim D Frank-Kamenetskii. Base-stacking and basepairing contributions into thermal stability of the DNA double helix. Nucleic acids research, 34(2):564– 574, 2006.

[101] Kristin A Eckert and Thomas A Kunkel. DNA polymerase fidelity and the polymerase chain reaction. Genome research, 1(1):17–24, 1991.

[102] James P Ferris. Montmorillonite-catalysed formation of RNA oligomers: the possible role of catalysis in the origins of life. Philosophical Transactions of the Royal Society B: Biological Sciences, 361(1474):1777– 1786, 2006.

[103] Jürgen Schnakenberg. Network theory of microscopic and macroscopic behavior of master equation systems. Reviews of Modern physics, 48(4):571, 1976.

[104] Jianshu Cao. Michaelis-Menten Equation and Detailed Balance in Enzymatic Networks. The Journal of Physical Chemistry B, 115(18):5493–5498, 2011.

